# Decreased Astrocytic CCL5 by MiR-324-5p Ameliorates Ischemic Stroke Injury via the CCR5/ERK/CREB Pathway

**DOI:** 10.1101/2024.09.03.610883

**Authors:** Jingxiu Li, Keyuan Gao, Lili Wang, Xinrui Wang, Kai Lian, Qingchen Ma, Xinmiao Ding, Yubing Wang, Chao Li, Zhiqin Gao, Chenxi Sun

## Abstract

Following ischemic stroke, Ccl5 mRNA expression increased, while miR-324-5p expression decreased in the peri-infarct cortex of middle cerebral artery occlusion (MCAO) mice. However, the roles of CCL5 and miR-324-5p in stroke remained largely unclear. Here, we show that inhibiting CCL5 using antibodies or miR-324-5p not only reduced infarct area and preserved neurological function in MCAO mice but also attenuated astrocyte reactivity and microglial activation, protected dendritic structures, and maintained spine density. In an astrocyte-neuron co-culture system after oxygen-glucose deprivation (OGD), inhibiting astrocytic CCL5 by antibody or miR-324-5p decreased neuronal apoptosis and preserved dendritic structure. Importantly, the suppression of CCL5 enhanced ERK/CREB pathway signaling both *in vivo* and *in vitro*. Consistent with these findings, the application of Maraviroc, a CCR5 antagonist, reduced infarct size, decreased neuronal apoptosis, and upregulated the ERK/CREB pathway in neurons treated with OGD. In conclusion, targeting the CCL5 pathway via miR-324-5p represents a promising therapeutic strategy for alleviating ischemic stroke damage through modulation of the neuronal CCR5/ERK/CREB pathway.

## Introduction

Stroke remains one of the leading causes of death and disability worldwide (Campbell et al. 2019). The most common subtype of stroke is ischemic stroke; however, its pathogenesis remains incompletely understood, and therapeutic options remain limited (H. Zhu et al. 2021). Astrocytes, the most abundant glial cells in the brain, serve diverse functions in the central nervous system (CNS), including contributing to the blood-brain barrier, synthesizing neurotransmitters, providing metabolic and trophic support, and modulating synaptic plasticity (Araque et al. 2014). Following a stroke, astrocytes are pivotal in mediating both the progression of neural injury and the ensuing protective responses, underscoring their critical and versatile roles in stroke pathophysiology.

The post-stroke period has been broadly divided into acute, subacute, and chronic phases (Dobkin and Carmichael 2016; Bernhardt et al. 2017). The acute phase and early subacute phase are critical periods of neuroplasticity (Biernaskie et al. 2004). During the acute phase of ischemic stroke, a large number of neurons undergo apoptosis, astrocytes undergo reactive astrogliosis and microglia become activated, the blood-brain barrier is disrupted, inflammatory mediators are upregulated, and immune cells infiltrate the injury site (Lambertsen et al. 2012; Kang et al. 2010). During the subacute phase, some neurons in the ischemic area undergo secondary death due to excitotoxicity or oxidative stress, while surviving neurons progressively regain function. Astrocytes remain reactive and microglia remain activated throughout this period, exerting both beneficial and detrimental effects. On the beneficial side, they phagocytose cellular debris, recruit immune cells, and regulate ionic homeostasis. On the detrimental side, however, they may exacerbate neuroinflammation and contribute to harmful processes such as excessive oxidative stress (Hamby et al. 2012; Cekanaviciute et al. 2014; Y. Zhu et al. 2002; Lin et al. 2023).

MicroRNAs (miRNAs) are small (∼22 nt) non-coding RNAs that bind to the 3’-UTR of target mRNAs, resulting in mRNA degradation or translational repression (J.-L. Mo et al. 2018). MicroRNAs are characterized by high sequence conservation, structural simplicity, and the capacity to cross the blood-brain barrier. These properties, together with their ease of synthesis as therapeutic agents, make them promising candidates for clinical diagnosis and therapeutic intervention (P. Sun et al. 2018). Under pathological conditions, miRNA expression profiles are dysregulated in the ischemic brain (Bhalala et al. 2013; S. Zhou et al. 2016), and modulation of astrocytic miRNA expression has been shown to confer neuroprotection against stroke-related inflammation, glutamate excitotoxicity, and glial scarring (Lekoubou et al. 2023; Li et al. 2021; Feigin et al. 2023; Nukovic et al. 2023). Our previous research demonstrated that miR-324-5p critically regulates C-C Motif Chemokine Ligand 5 (CCL5, also known as RANTES: Regulated on Activation, Normal T Cell Expressed and Secreted) expression, playing an essential role in synaptic formation by inhibiting CCL5 secretion from astrocytes (C. Sun et al. 2019). However, in the context of stroke, the impact of astrocytic CCL5 regulation via miR-324-5p on neurological function and downstream neuronal signaling pathways remains largely unknown.

Chemokines, including CCL5, are a subclass of pro-inflammatory cytokines with chemoattractant properties that serve as important regulators of both peripheral and central immune responses (Baggiolini 1998; Réaux-Le Goazigo et al. 2013; Trettel et al. 2020). CCL5 is produced by a variety of cell types, including T lymphocytes, platelets, endothelial cells, smooth muscle cells, and glial cells (Julián-Villaverde et al. 2022). Whether circulating CCL5 levels are elevated or reduced following ischemic stroke remains a matter of debate, with conflicting findings across studies (Julián-Villaverde et al. 2022). Some studies have reported elevated CCL5 levels in both blood and cerebrospinal fluid (Pawluk et al. 2023; Tokami et al. 2013; Montecucco et al. 2010; Q. Kong et al. 2020), while others have found no significant differences in CCL5 levels compared to controls or among ischemic stroke patients across different time points (García-Berrocoso et al. 2014; Julián-Villaverde et al. 2022; Zaremba et al., n.d.).

In the CNS, CCL5 and its receptors serve multiple functions, including promoting neuroinflammation, modulating synaptic activity, and conferring neuroprotection against various neurotoxins (Lanfranco et al. 2018). Studies using CCL5-knockout mice have demonstrated a reduction in ischemic lesion volume and significantly reduced blood-brain barrier disruption, along with decreased leukocyte and platelet adhesion (Terao et al. 2008). Additionally, administration of a CCL5 inhibitor in experimental stroke models has been shown to decrease infarct size and improve neurological outcomes post-stroke (Fan et al. 2016). Conversely, elevated CCL5 levels may confer neuroprotection through mechanisms such as vasodilation, inhibition of platelet aggregation, and promotion of angiogenesis (Julián-Villaverde et al. 2022). Furthermore, CCL5 has been reported to upregulate the expression of neurotrophic factors such as brain-derived neurotrophic factor and epidermal growth factor (Tokami et al. 2013). Overexpression of CCL5 in wild-type mice has been associated with enhanced learning and memory performance, as well as increased hippocampal neuronal activity and connectivity (Ajoy et al. 2021). These contradictory findings underscore the complexity of CCL5 function in the injured brain and highlight the need for more detailed investigations into the regulatory mechanisms governing CCL5 following stroke.

CCL5 signals through multiple receptors, notably C-C Motif Chemokine Receptor 5 (CCR5), CCR1, CCR3, and GPR75 (Ajoy et al. 2021). CCR5, a predominant receptor for CCL5, is a seven-transmembrane G-protein–coupled receptor that plays a crucial role in regulating pro-inflammatory responses by modulating immune cell behavior, survival, and tissue retention (Kohlmeier et al. 2011). CCR5 is also expressed in non-immune cells such as astrocytes, microglia, and neurons, where it influences neuronal survival and differentiation (Sorce et al. 2010). CCR5 mRNA and protein levels were found to be increased in the brain, spinal cord, and blood plasma at 22 hours following middle cerebral artery occlusion (MCAO) in mice (Yan et al. 2021). Stroke induces an upregulation of CCR5 expression in neurons and a simultaneous downregulation in microglia, indicating complex, cell-type-specific changes in CCR5 signaling during the recovery phase (Joy et al. 2019).

In this study, building upon our earlier work on miR-324-5p-mediated regulation of astrocytic CCL5 expression, we investigated the neuroprotective effects of CCL5 inhibition via neutralizing antibodies or miR-324-5p *in vivo* using the MCAO model. Subsequently, we examined the role of reduced astrocytic CCL5 expression in inhibiting neuronal apoptosis and preserving dendritic structure using the oxygen-glucose deprivation (OGD) model *in vitro*. Finally, we explored the interaction between astrocytic miR-324-5p/CCL5 signaling and the neuronal CCR5/ERK/CREB pathway both in the peri-infarct cortex of MCAO mice and in primary cortical neurons following OGD.

## Results

To characterize regional differences in gene expression and protein levels following ischemic stroke, we sampled three cortical regions in MCAO mice: the Ipsilateral Core (IC), Ipsilateral Penumbra (IP), and Contralateral Penumbra (CP), as illustrated in Figure 1B. The IC region undergoes irreversible neuronal death due to severe oxygen deprivation, while the IP region sustains comparatively less damage and represents the principal target for therapeutic intervention, where neurons may either die or recover following ischemia-reperfusion injury. We therefore focused our analyses on the IP region.

**Figure 1.**
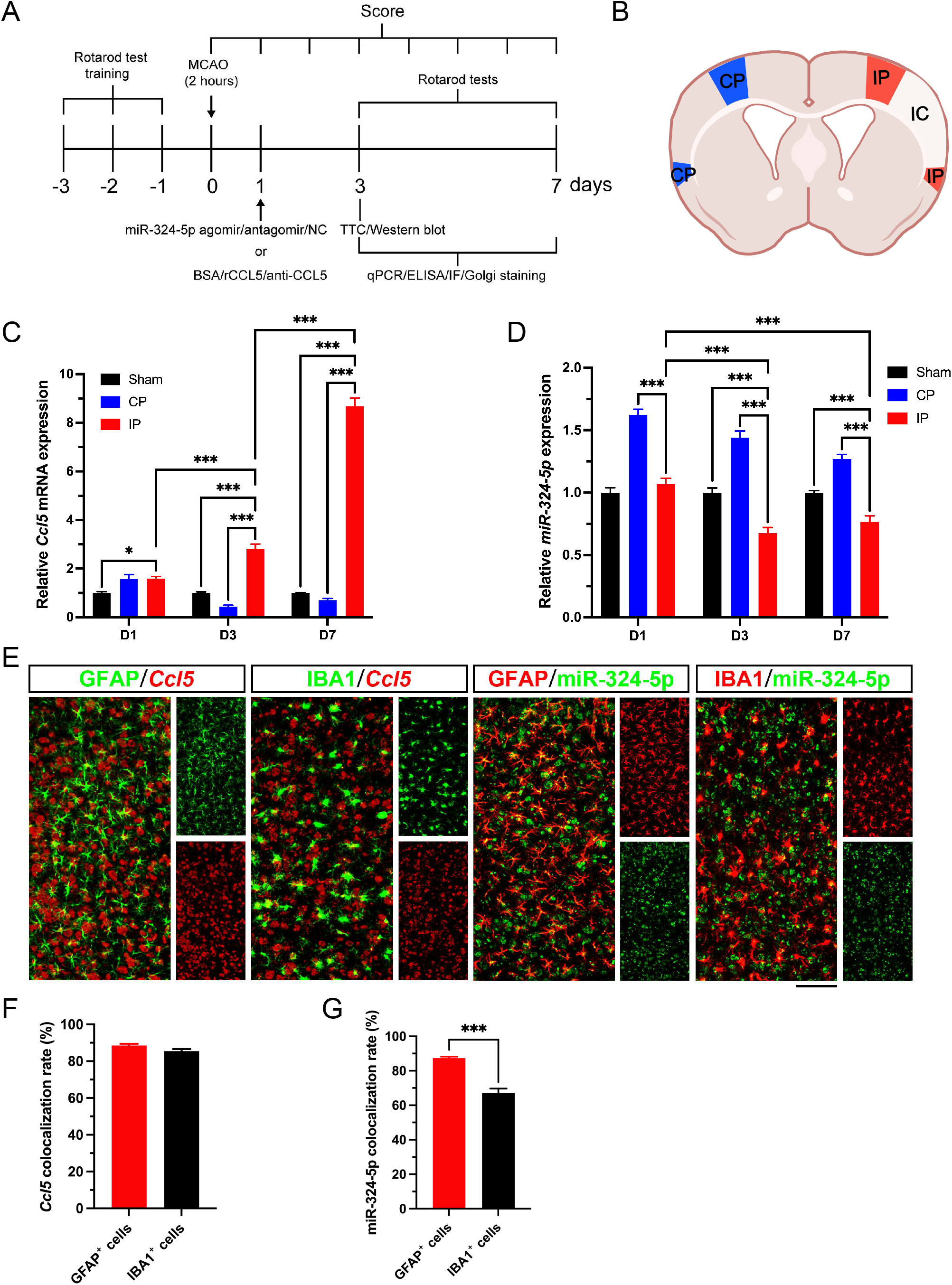
Ccl5 mRNA expression is upregulated and miR-324-5p is downregulated in the peri-infarct cortex of MCAO mice. (A) Illustration of the animal experimental protocol. (B) Schematic diagram of the brain indicating IC, IP, and CP cortical regions analyzed in this study. (C and D) qRT-PCR analysis of Ccl5 mRNA (C) and miR-324-5p (D) expression in the IP and CP regions on D1, D3, and D7 after MCAO, and in sham mice (*n*=6). (E) Representative images of combined FISH for Ccl5 mRNA or miR-324-5p and immunofluorescence for GFAP or IBA1 in the IP region on D3 after MCAO. Scale bar: 100 μm. (F and G) Colocalization rate of Ccl5 mRNA probe (F) or miR-324-5p probe (G) with GFAP-positive astrocytes or IBA1-positive microglia (*n*=10). Statistical comparisons in C and D were performed by two-way ANOVA with Tukey’s post-hoc test. In F and G, comparisons between astrocytes and microglia were made by unpaired Student’s *t*-test.

As depicted in Figure 1C, quantitative real-time PCR (qRT-PCR) results indicated that Ccl5 mRNA levels in the IP region were significantly increased compared with sham controls on D1, D3, and D7. Compared with the CP region, Ccl5 mRNA levels in the IP region were significantly higher at D3 and D7. Furthermore, Ccl5 mRNA levels continuously increased from D1 to D7 in the IP region after MCAO.

As shown in Figure 1D, qRT-PCR results demonstrated that miR-324-5p levels in the IP region were significantly lower than sham controls on D3 and D7. Compared with the CP region, miR-324-5p levels in the IP region were significantly lower at D1, D3, and D7. Furthermore, miR-324-5p levels in the IP region were markedly decreased at D3 and D7 relative to D1, although no significant differences were observed between D3 and D7.

Taken together, the results in Figure 1C and 1D show that Ccl5 mRNA levels in the IP region were significantly elevated from D1 onwards compared with sham. At D1, miR-324-5p levels in the IP region did not differ significantly from sham, although they were significantly lower than in the CP region. On D3 and D7, miR-324-5p levels in the IP region were significantly downregulated compared with both sham and CP groups, which may have diminished its inhibitory effect on Ccl5 expression, and contributed to the continuous increase in Ccl5 mRNA levels in the IP region.

CCL5 is a well-known pro-inflammatory factor that, in the brain, is mainly produced and released by microglia and astrocytes. Following brain injury or inflammation, CCL5 has been reported to serve as an indicator of the inflammatory response. To confirm the miR-324-5p/CCL5 axis in glial cells after ischemic stroke, fluorescence *in situ* hybridization (FISH) was performed to detect the *in situ* expression of Ccl5 mRNA and miR-324-5p at cellular levels in the peri-infarct cortex of MCAO mice at D3. As shown in Figure 1E-F, Ccl5 mRNA was detected at both astrocytes and microglia, with a similar Ccl5 colocalization rate between the two cell types. In contrast, miR-324-5p showed a significantly higher colocalization rate in astrocytes than in microglia (Figure 1G). Thus, both astrocytes and microglia expressed Ccl5 and miR-324-5p in the peri-infarct region after MCAO; given the higher miR-324-5p expression in astrocytes, the miR-324-5p/CCL5 axis may exert a more pronounced regulatory effect on astrocytes.

To investigate the effects of the miR-324-5p/CCL5 axis in MCAO mice, recombinant mouse CCL5 protein (rCCL5), CCL5 antibody (anti-CCL5), or bovine serum albumin (BSA) was injected into the ipsilateral cortex at 24 h post-surgery (Figure 1A). Structural and functional analyses were subsequently performed from D3 to D7. In addition, the CCR5 antagonist Maraviroc (MVC, also known as UK-427857) was administered intraperitoneally at 4, 24, and 48 h post-surgery to evaluate its effect on recovery after ischemic stroke.

CCL5 protein levels in cortical tissue from MCAO mice were measured using ELISA on D3 and D7 (Figure 2A). In the BSA control group on D3 and D7, CCL5 concentrations in the IP region were notably higher than in the CP region, and concentrations on D7 were significantly higher than those on D3. These results were consistent with the elevated Ccl5 mRNA levels detected by qRT-PCR in MCAO mice. In the IP regions on D3 and D7, significantly reduced CCL5 concentrations were observed in the anti-CCL5 group compared with the BSA control group, whereas CCL5 concentrations in the rCCL5 group were notably higher compared to other groups. These results suggest that injection of CCL5 antibody into the peri-infarct cortex significantly decreased CCL5 concentrations from D3 to D7, whereas rCCL5 injection significantly increased CCL5 protein concentrations in the IP region from D3 to D7. Continuous CCR5 blockade with MVC also decreased CCL5 concentrations in the IP region compared with the BSA controls at D3 and D7.

**Figure 2.**
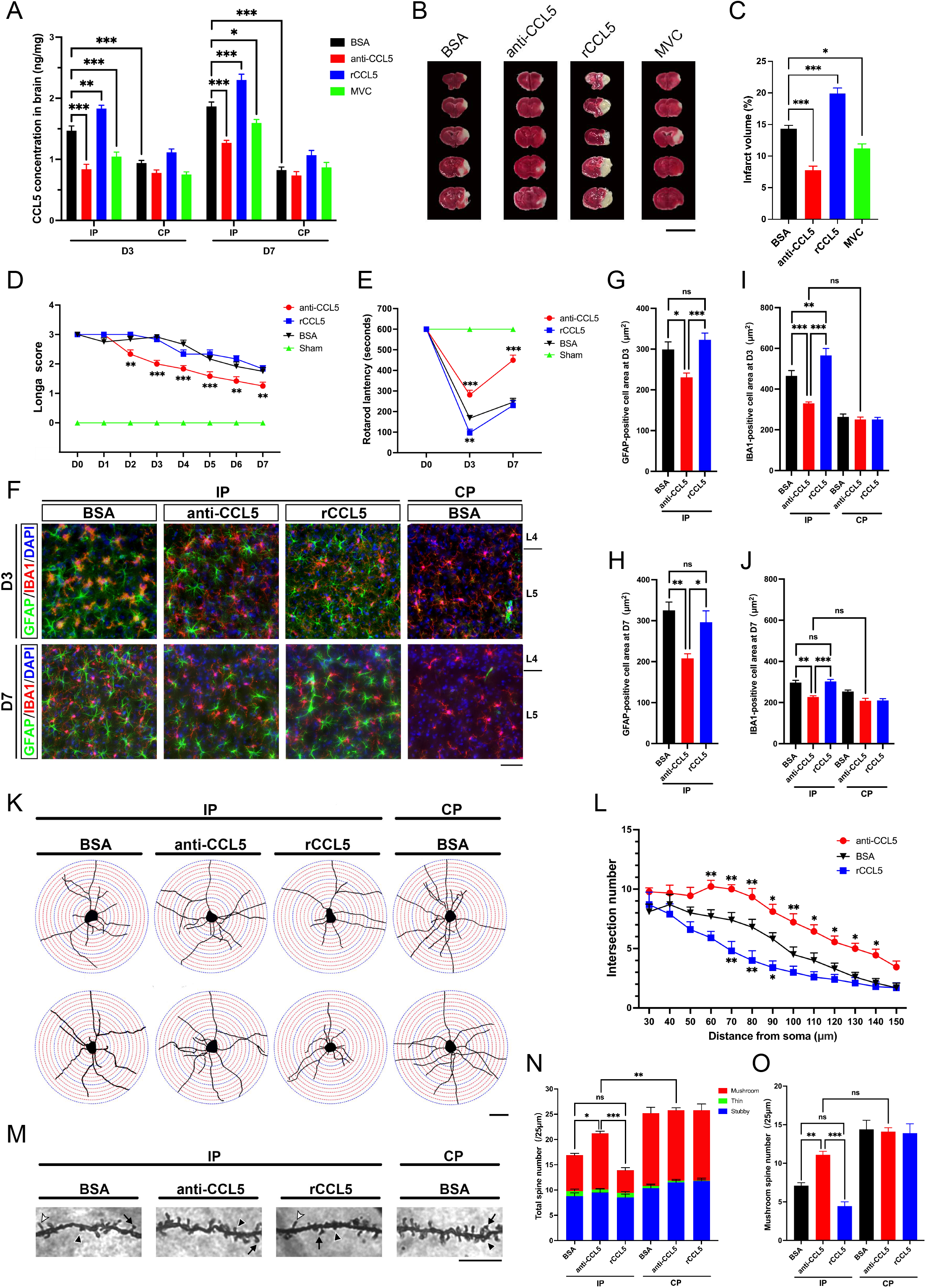
CCL5 inhibition by intracerebral antibody delivery and CCR5 blockade attenuate ischemic stroke injury. (A) ELISA quantification of CCL5 concentrations in the IP and CP regions of BSA-, CCL5 antibody-, rCCL5-, and MVC-treated mice on D3 and D7 after MCAO (*n*=6). (B and C) Representative TTC-stained brain sections (B) and infarct volume (C) of BSA-, CCL5 antibody-, rCCL5-, and MVC-treated mice on D3 after MCAO (*n*=6). Scale bar: 1cm. (D) Longa scores in BSA-, CCL5 antibody-, and rCCL5-treated mice, and sham controls, from D0 to D7 after MCAO (*n*=12). (E) Rotarod performance of BSA-, CCL5 antibody-, and rCCL5-treated mice, and sham controls, on D0, D3, and D7 after MCAO (*n*=7). (F) Representative immunofluorescence images of GFAP and IBA1 co-labeling in the IP cortex of BSA-, CCL5 antibody-, and rCCL5-treated mice, and in the CP cortex of BSA-treated mice, on D3 and D7 after MCAO. Scale bar: 50 μm. (G and H) Surface area of GFAP-positive astrocytes in the IP region of BSA-, CCL5 antibody-, and rCCL5-treated mice on D3 (G) and D7 (H) after MCAO (*n*=10). (I and J) Surface area of IBA1-positive microglia in the IP region of BSA-, CCL5 antibody-, and rCCL5-treated mice on D3 (I) and D7 (J) after MCAO (*n*=10). (K) Representative tracings of Golgi-stained cortical neurons illustrating dendritic morphology in the IP cortex of BSA-, CCL5 antibody-, and rCCL5-treated mice, and in the CP cortex of BSA-treated mice, on D3 after MCAO. Scale bar: 50 μm. (L) Sholl analysis of the dendritic branch complexity in the IP region of BSA-, CCL5 antibody-, and rCCL5-treated mice on D3 after MCAO (*n*=10). (M) Representative images of basal dendritic spines in the IP cortex of BSA-, CCL5 antibody-, and rCCL5-treated mice, and in the CP cortex of BSA-treated mice, on D7 after MCAO. Arrows, mushroom spines; black arrowheads, stubby spines; white arrowheads, thin spines. Scale bar: 10 μm. (N and O) Total spine density (N) and mushroom spine density (O) in the IP and CP regions of BSA-, CCL5 antibody-, and rCCL5-treated mice on D7 after MCAO (*n*=10). Stacked columns represent total spine density, with colored segments indicating the proportions of mushroom, stubby, and thin spines. Statistical comparisons in A, D, E, and L were performed by two-way ANOVA with Tukey’s post-hoc test; all other panels by one-way ANOVA with Tukey’s post-hoc test.

The 2,3,5-triphenyltetrazolium chloride (TTC) results indicated that at D3, the infarct area in the anti-CCL5 group significantly decreased compared with the BSA control (Figure 2B-C). In contrast, following rCCL5 injection, the infarct area significantly increased, reaching approximately 1.4 ± 0.1 times that of the BSA group. Furthermore, after MVC injection, the infarct area in MCAO mice significantly decreased compared with the BSA controls. These results demonstrate that reducing cortical CCL5 protein expression or blocking the CCR5 receptor after MCAO significantly reduced neuronal death after ischemic injury, thereby exerting a protective effect against ischemic brain damage. Conversely, increasing CCL5 protein levels significantly increased the infarct area after ischemic injury, indicating its role in exacerbating ischemic cortical damage.

Following injection of the CCL5 antibody, MCAO mice exhibited significantly lower Longa scores compared with the BSA group from D2 to D7 (Figure 2D). No statistically significant difference in the score was observed between the BSA group and the rCCL5 group. Sham mice recovered to a score of 0 by 24 hours post-operation and maintained this score through D7, indicating that, aside from occlusion, the surgery did not significantly affect the motor behavior of the mice. These results indicate that reducing CCL5 expression in the ipsilateral cortex of MCAO mice significantly protects motor function after ischemic brain injury.

All mice successfully completed the rotarod test before MCAO surgery and remained on the rod without falling. At both D3 and D7, mice in the anti-CCL5 group showed significantly longer latency-to-fall on the rotarod compared with BSA control groups (Figure 2E). In contrast, significantly shorter latency-to-fall on the rotarod were detected in the rCCL5 group compared with the BSA control group at D3. By D7, no statistically significant differences were detected between the two groups. Sham mice remained on the rotarod without falling at both D3 and D7, indicating that aside from occlusion, the surgical procedure did not significantly affect the rotarod performance of the mice. These findings indicate that reducing CCL5 expression in the ipsilateral cortex improved neurobehavioral recovery in MCAO mice.

GFAP/IBA1 immunofluorescence staining was employed to detect and distinguish reactive astrocytes and activated microglia within the IP cortex (Figure 2F). Statistical analysis indicated that at both D3 and D7, the surface areas of GFAP-positive astrocytes and IBA1-positive microglia were significantly reduced in the anti-CCL5 group (Figure 2G-J). Moreover, in the anti-CCL5 group, no statistical differences were observed in microglial surface area between the IP and CP regions (Figure 2I-J). Conversely, astrocyte surface area in the rCCL5 group showed no significant difference compared with the BSA group (Figure 2G-H). Notably, at D3, microglial surface area in the rCCL5 group was significantly larger than in the BSA control group (Figure 2I), whereas by D7, microglial surface areas in both groups decreased to comparable levels (Figure 2J). No significant differences were detected in the density of reactive astrocytes or activated microglia in the IP region among the BSA, anti-CCL5, and rCCL5 groups at both D3 and D7 (Supplementary Figure S1A-D). These results suggest that reducing CCL5 concentration in the injured cortex of MCAO mice suppresses astrocyte reactivity and microglial activation, potentially reducing brain inflammation after ischemic injury.

Sholl analysis performed after Golgi staining revealed that at D3 and D7, the dendrite intersections of cortical neurons in the IP region of the anti-CCL5 group were significantly more than those in the BSA control group (Figure 2K-L, Supplementary Figure S1E). At D3, dendritic branch complexity of cortical neurons in the IP region of the anti-CCL5 group was significantly higher within the 60-140 μm range from the soma compared with the BSA control group. In the IP region of the rCCL5 group, dendritic branch complexity was significantly lower compared with the BSA group within the 70-90 μm range from the soma. No significant differences were found in dendritic branch complexity in the CP cortex among the three groups (Supplementary Figure S1F). These results indicate that inhibiting CCL5 expression in the ischemic cortex significantly protects neuronal dendritic structure, while excess CCL5 expression further compromises the dendritic structure in cortical neurons.

We next analyzed differences in dendritic spine density and morphology among the groups. In the IP region of the anti-CCL5 group at D3 and D7, both the total dendritic spine density and the density of mature mushroom-type dendritic spines were significantly higher compared with the rCCL5 and BSA control groups, while the densities of thin-type and stubby-type immature dendritic spines showed no significant differences across groups (Figure 2M-O, Supplementary Figure S1G-H). No statistical differences were observed in total dendritic spine density or in the proportions of different spine morphologies between the rCCL5 group and the BSA group in the IP region. Likewise, no statistical differences were observed in the total dendritic spine density or in the proportions of the three different morphological types of dendritic spines in the CP cortical neurons among the groups. These results suggest that inhibiting CCL5 expression in the injured cortex after ischemic stroke has a significant protective effect on dendritic spine density, particularly of the mature mushroom-type dendritic spines.

Having examined the effects of overexpressing or inhibiting CCL5 protein on neurological function, glial reactivity, and neuronal structure following ischemic stroke, we next administered *in situ* injections to the ipsilateral cortex of MCAO mice to modulate miR-324-5p levels. ELISA results revealed that in the negative control (NC) group on D3 and D7, CCL5 concentrations were notably higher in the IP regions than in CP regions (Figure 3A). Compared with the NC control, CCL5 protein concentrations in the IP regions of the miR-324-5p agomir group decreased significantly on D3 and D7. Conversely, the miR-324-5p antagomir group exhibited a significant increase in CCL5 concentration in the IP region at D3 and D7. These findings indicate that miR-324-5p agomir injection significantly reduced CCL5 expression in the ischemic cortex, whereas antagomir injection markedly increased it.

**Figure 3.**
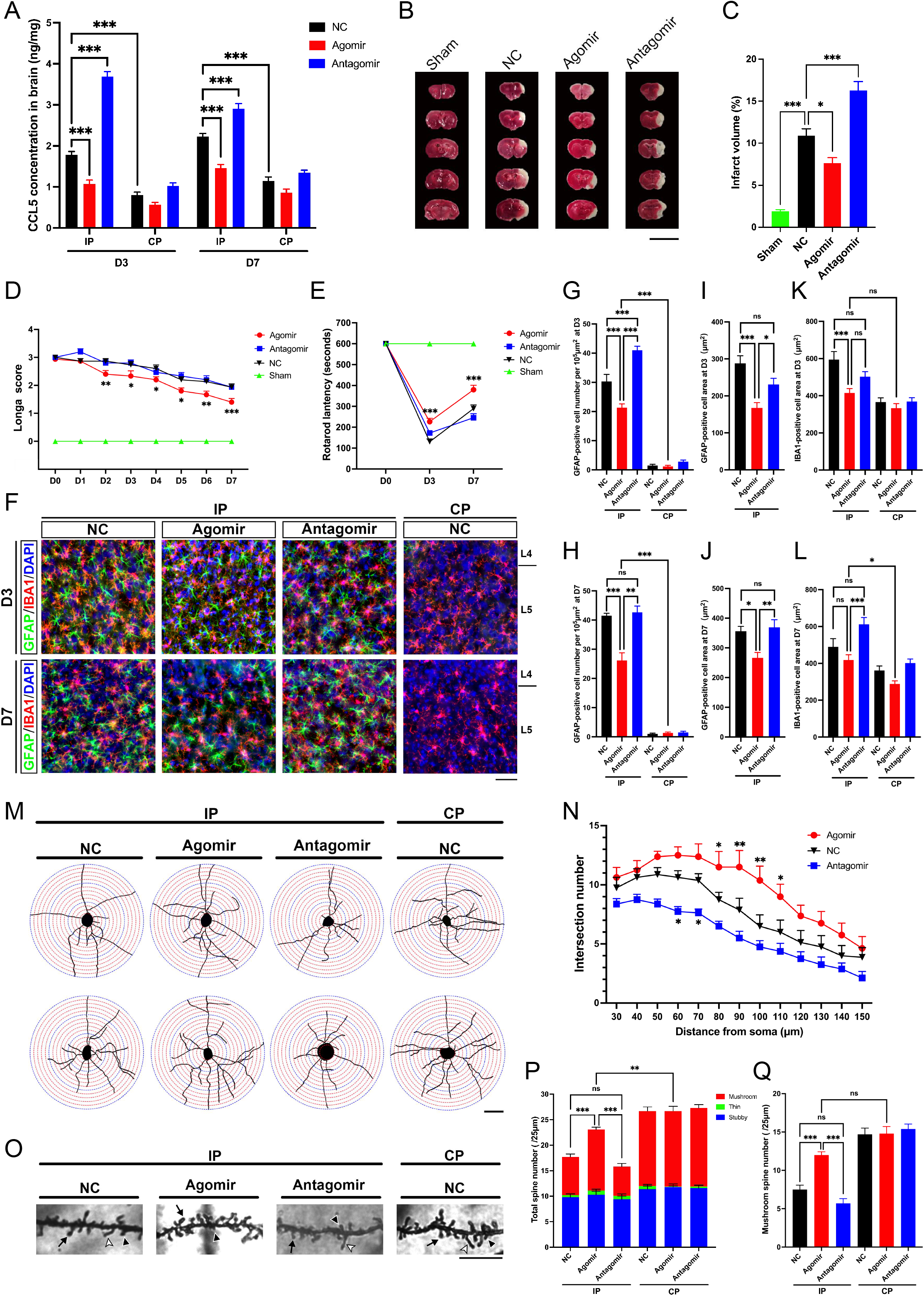
Intracerebral miR-324-5p agomir delivery reduces ischemic stroke injury. (A) ELISA quantification of CCL5 concentrations in the IP and CP regions of NC agomir-, miR-324-5p agomir-, and miR-324-5p antagomir-treated mice on D3 and D7 after MCAO (*n*=6). (B and C) Representative TTC-stained brain sections (B) and infarct volume (C) in sham, NC agomir-, miR-324-5p agomir-, and miR-324-5p antagomir-treated mice on D3 after MCAO (*n*=6). Scale bar: 1cm. (D) Longa scores in NC agomir-, miR-324-5p agomir-, and miR-324-5p antagomir-treated mice, and sham controls, from D0 to D7 after MCAO (*n*=15). (E) Rotarod performance of NC agomir-, miR-324-5p agomir-, and miR-324-5p antagomir-treated mice, and sham controls, on D0, D3, and D7 after MCAO (*n*=7). (F) Representative immunofluorescence images of GFAP and IBA1 co-labeling in the IP cortex of NC agomir-, miR-324-5p agomir-, and miR-324-5p antagomir-treated mice, and in the CP cortex of NC agomir-treated mice, on D3 and D7 after MCAO. Scale bar: 50 μm. (G and H) Density of GFAP-positive astrocytes (cells per 10^5^ μm^2^) in the IP and CP regions of NC agomir-, miR-324-5p agomir-, and miR-324-5p antagomir-treated mice on D3 (G) and D7 (H) after MCAO (*n*=6). (I and J) Surface area of GFAP-positive astrocytes in the IP region of NC agomir-, miR-324-5p agomir-, and miR-324-5p antagomir-treated mice on D3 (I) and D7 (J) after MCAO (*n*=10). (K and L) Surface area of IBA1-positive microglia in the IP and CP regions of NC agomir-,miR-324-5p agomir-, and miR-324-5p antagomir-treated mice on D3 (K) and D7 (L) after MCAO (*n*=10). (M) Representative tracings of Golgi-stained cortical neurons illustrating dendritic morphology in the IP cortex of NC agomir-, miR-324-5p agomir-, and miR-324-5p antagomir-treated mice, and in the CP cortex of NC agomir-treated mice, on D3 after MCAO. Scale bar: 50 μm. (N) Sholl analysis of dendritic branch complexity in the IP region of NC agomir-, miR-324-5p agomir-, and miR-324-5p antagomir-treated mice on D3 after MCAO (*n*=8). (O) Representative images of basal dendritic spines in the IP cortex of NC agomir-,miR-324-5p agomir-, and miR-324-5p antagomir-treated mice, and in the CP cortex of NC agomir-treated mice, on D7 after MCAO. Arrows, mushroom spines; black arrowheads, stubby spines; white arrowheads, thin spines. Scale bar: 10 μm. (P and Q) Total spine density (P) and mushroom spine density (Q) in the IP and CP regions of NC agomir-, miR-324-5p agomir-, and miR-324-5p antagomir-treated mice on D7 after MCAO (*n*=10). Stacked columns represent total spine density, with colored segments indicating the proportions of mushroom, stubby, and thin spines. Statistical comparisons in A, D, E, and N were performed by two-way ANOVA with Tukey’s post-hoc test; all other panels by one-way ANOVA with Tukey’s post-hoc test.

The TTC results revealed that the infarct area was significantly smaller in the miR-324-5p agomir group compared with the NC agomir group at D3 (Figure 3B-C). Conversely, after miR-324-5p antagomir injection, the infarct area significantly increased. The infarct area in sham mice was less than 2.5% of the total brain area, indicating that, aside from occlusion, the surgery did not cause substantial infarction. These results demonstrate that miR-324-5p overexpression reduces cortical neuronal death and exerts a neuroprotective effect following ischemic stroke, while its inhibition exacerbates neuronal death and increases infarct area.

The Longa score results indicated that miR-324-5p agomir-treated mice consistently exhibited significantly lower scores than the NC agomir control group from D2 to D7 (Figure 3D), while no significant differences were observed between the antagomir and NC control groups. These data suggest that miR-324-5p overexpression in the IP regions significantly enhances neurological function following ischemic stroke.

The rotarod tests indicated that on D3 and D7, mice in the agomir group exhibited significantly longer latency-to-fall compared with NC groups (Figure 3E), whereas no statistically significant differences were found between the antagomir and NC groups. These results suggest that miR-324-5p promotes sensorimotor recovery following ischemic brain injury.

Statistical analysis revealed that following the injection of miR-324-5p agomir, the density of GFAP-positive reactive astrocytes in the IP region significantly decreased on D3 and D7, compared with both the NC control and the antagomir groups (Figure 3G-H). The density of reactive astrocytes in the IP region in the antagomir group was significantly higher than in the NC control group on D3. However, due to an increase in reactive astrocytes in the NC group by D7, no statistical differences were observed between the antagomir group and the NC group at this time point. Furthermore, the surface area of reactive astrocytes in the IP region was smaller in the agomir group compared with the NC group on both D3 and D7, while no statistical differences were observed between the antagomir group and the NC group (Figure 3I-J).

On D3, microglial surface area in the agomir group was significantly smaller than in the NC group (Figure 3K). By D7, due to decreased microglial surface area in the NC group and increased microglial surface area in the antagomir group, no significant difference between the agomir group and the NC group was detected, but microglial surface area in the agomir group remained significantly smaller than in the antagomir group (Figure 3L). On both D3 and D7, no statistically significant differences were observed in activated microglial surface area between the antagomir group and the NC group in the IP region. Moreover, no significant differences were detected in the density of activated microglia between the agomir and NC groups in the IP regions on both D3 and D7 (Supplementary Figure S2A-B). These results indicate that miR-324-5p overexpression following ischemic stroke significantly inhibited the reactive transformation and proliferation of astrocytes in the ipsilateral cortex, and also reduced microglial activation, as evidenced by the reduction in microglial surface area on D3.

Sholl analysis of Golgi-stained cortical neurons in the IP region revealed that at D3 and D7, the dendrite intersections of cortical neurons in the IP region of the miR-324-5p agomir group were significantly more than those in the NC control group (Figure 3M-N, Supplementary Figure S2C). At D3, significantly more dendritic intersections were observed within 80-110 μm from the soma in the agomir group compared with the NC group. Conversely, in the antagomir group, dendritic intersections were significantly fewer than those in the NC group within 60-70 μm from the soma. No significant differences were found in dendritic branch complexity in the CP cortex among the three groups (Supplementary Figure S2D). These findings suggest that miR-324-5p overexpression after ischemic brain injury preserves neuronal dendritic structures, potentially facilitating functional recovery following ischemic stroke.

Analysis of dendritic spine density and morphology in the IP region demonstrated that on D3 and D7, both the total dendritic spine density and the density of mushroom-type mature dendritic spines in the agomir group were significantly higher than in both the NC control and the antagomir groups (Figure 3O-Q, Supplementary Figure S2E-F). No significant differences in total dendritic spine density and mushroom-type mature dendritic spine density were observed between the antagomir group and the NC group at D3 and D7. These results underscore the significant protective effects of increasing miR-324-5p expression on neuronal dendritic spine density post-ischemic stroke, particularly for mature mushroom-type dendritic spines.

To examine the effects of astrocytic CCL5 on neuronal survival and dendritic integrity under ischemic conditions, we established an *in vitro* co-culture model of primary cortical astrocytes and neurons, as illustrated in Figure 4A. Following 1 h OGD treatment, the co-culture media were supplemented with BSA, CCL5 antibody, or rCCL5 to examine the effects of modulating CCL5 expression on neuronal apoptosis and dendritic structure. ELISA was performed to measure CCL5 concentration in the co-cultured medium at D3 post-OGD (Figure 4B). Results showed that the CCL5 concentration in the un-OGD-treated control group was approximately 45.0 ± 5.4 pg/mL. Following OGD treatment, CCL5 concentration significantly increased in the BSA control group, reaching 145.0 ± 7.6 pg/mL. The addition of CCL5 antibody significantly decreased CCL5 concentrations in the OGD-treated co-cultured medium compared with the BSA control, reducing it to 48.3 ± 6.8 pg/mL. Conversely, the supplementation with rCCL5 led to a significant increase in CCL5 concentration, reaching 829.6 ± 37.1 pg/mL. These findings indicate that astrocytic CCL5 expression significantly increased after OGD treatment. The introduction of CCL5 antibody substantially reduced CCL5 concentrations in the extracellular milieu of the co-culture system, whereas exogenous rCCL5 substantially elevated CCL5 concentration. Moreover, when MVC was added to the co-culture media post-OGD, CCL5 concentration was significantly reduced. This result suggests that blocking astrocytic CCR5 reduces CCL5 production, thereby downregulating its downstream effects following OGD or ischemic injury.

**Figure 4.**
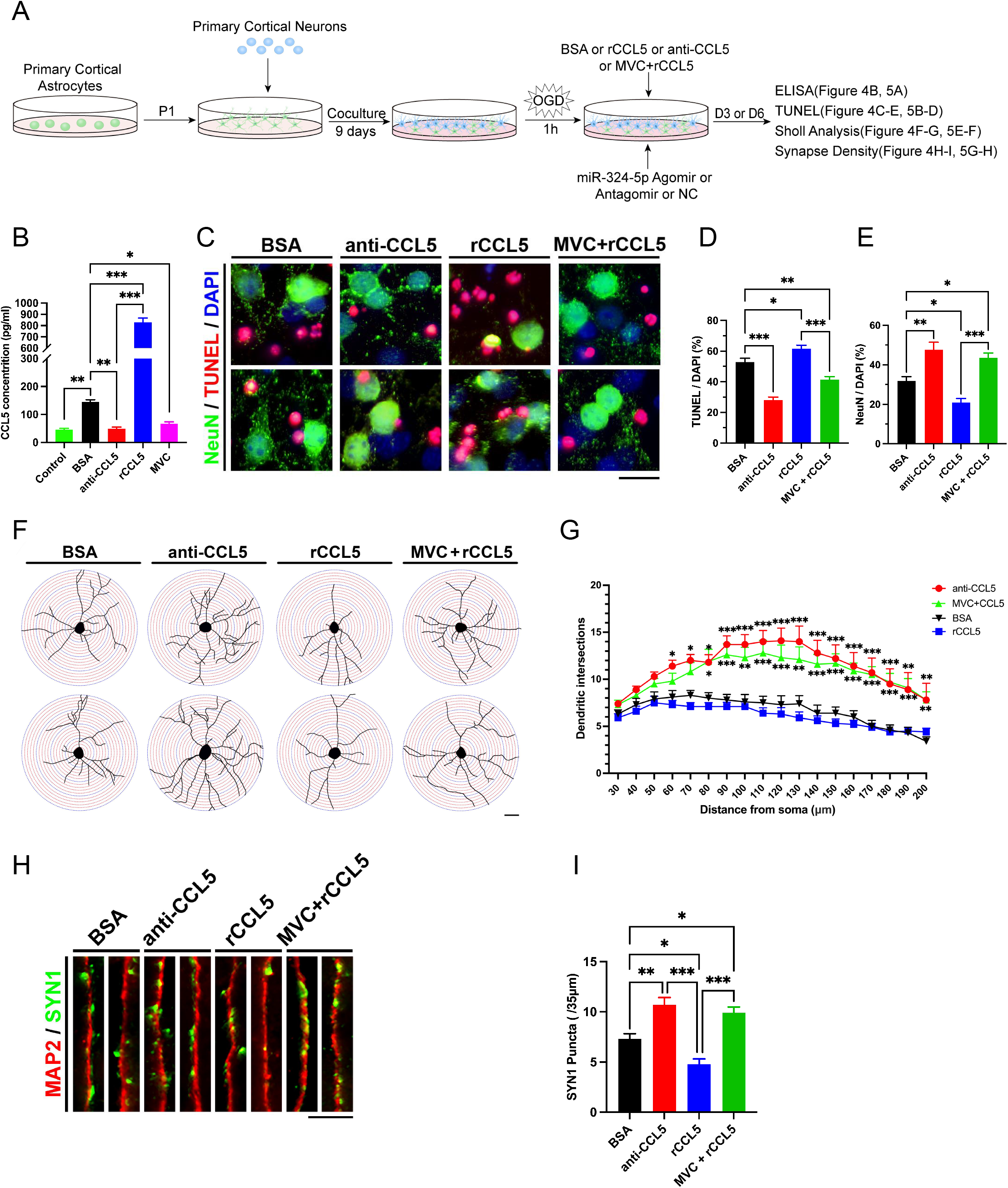
CCL5 inhibition and CCR5 blockade protect cortical neurons from OGD injury in astrocyte-neuron co-cultures. (A) Schematic of the astrocyte-neuron co-culture and OGD experimental design. (B) ELISA quantification of CCL5 concentrations in the medium of BSA-, CCL5 antibody-, rCCL5-, and MVC+rCCL5-treated co-cultures, and in un-OGD-treated controls (*n*=5). (C) Representative immunofluorescence images of NeuN and TUNEL co-labeling. Scale bar: 20 μm. (D and E) Proportions of apoptotic cells (D) and NeuN-positive neurons (E) in BSA-, CCL5 antibody-, rCCL5-, and MVC+rCCL5-treated co-cultures (*n*=10). (F and G) Representative tracings of MAP2-stained neurons (F) and Sholl analysis (G) of neuronal dendritic morphology in BSA-, CCL5 antibody-, rCCL5-, and MVC+rCCL5-treated co-cultures (*n*=10). Scale bar: 50 μm. (H and I) Representative immunofluorescence images of MAP2 and SYN1 co-labeling (H) and synapse puncta density (I) in BSA-, CCL5 antibody-, rCCL5-, and MVC+rCCL5-treated co-cultured neurons (*n*=10). Scale bar: 10 μm. Statistical comparisons in G were performed by two-way ANOVA with Tukey’s post-hoc test; asterisks positioned above indicate anti-CCL5 vs. BSA comparisons, and those positioned below indicate MVC+rCCL5 vs. BSA comparisons. All other panels were analyzed by one-way ANOVA with Tukey’s post-hoc test.

TUNEL staining indicated significantly fewer apoptotic cells in the anti-CCL5 group compared with the BSA control group on D3 after OGD (Figure 4C-D). Furthermore, the surviving NeuN-positive neurons was significantly more than in the BSA control (Figure 4E). In the rCCL5 group, the apoptotic cell ratio was significantly higher than in the BSA group, with fewer NeuN-positive neurons observed in the rCCL5 group. Simultaneous addition of rCCL5 and MVC (MVC+rCCL5) resulted in significantly fewer apoptotic cells compared with both the rCCL5 group and the BSA control. NeuN-positive neurons in the MVC+rCCL5 group were significantly more than in both the rCCL5 group and the BSA group. These results demonstrate that reducing CCL5 levels in the extracellular milieu of the co-culture system after OGD treatment, or inhibiting the CCR5 receptor, both exert a protective effect on neuronal survival following OGD injury. Conversely, excessive CCL5 leads to increased neuronal death after OGD.

Sholl analysis revealed that at D3 and D6 post-OGD, the dendrite intersections of neurons in the anti-CCL5 group were significantly more than those in the BSA control group (Figure 4F-G, Supplementary Figure S3A). At D6 post-OGD, dendrite field complexity in neurons treated with CCL5 antibody was significantly higher compared with the BSA control at distances of 60-200 μm from the soma. Compared with the BSA group, neurons in the MVC+rCCL5 group showed significantly increased dendrite intersections at distances of 80-200 μm from the soma. No significant difference was observed between rCCL5 and BSA groups. These results indicate that reducing CCL5 levels in the extracellular milieu of the co-culture system or inhibiting the CCR5 receptor significantly protects the dendritic structure of neurons after OGD injury.

MAP2/SYN1 immunofluorescence staining revealed that at D6 post-OGD treatment, neurons in the CCL5 antibody group exhibited a significantly higher synaptic density than those in the BSA control groups (Figure 4H-I). No significant differences were observed in synaptic density between the rCCL5 and BSA control groups. Notably, neuronal density in the MVC+rCCL5 group was markedly higher compared with the rCCL5 and BSA groups. These findings suggest that reducing CCL5 expression or inhibiting the CCR5 receptor can significantly preserve neuronal synaptic density, while excessive CCL5 exacerbated synapse loss following OGD injury.

Using the same co-culture system (Figure 4A), we next transfected NC agomir, miR-324-5p agomir, or miR-324-5p antagomir into the co-cultures to examine whether modulating miR-324-5p expression could recapitulate the effects of CCL5 inhibition on neuronal survival and dendritic integrity following OGD. On D3 and D6 post-OGD, the transfection efficiency of agomirs in the co-culture system reached approximately 60% (Supplementary Figure S3B). At D3 post-OGD, ELISA was used to measure CCL5 concentrations in the co-culture medium (Figure 5A). Results indicated that CCL5 levels in the agomir-transfected group were significantly lower than in both the NC group and antagomir groups. Conversely, CCL5 concentrations were significantly elevated in the antagomir-transfected group compared with the NC group. These findings demonstrate that agomir transfection significantly reduced CCL5 expression in the co-culture medium, whereas antagomir transfection significantly increased CCL5 expression.

**Figure 5.**
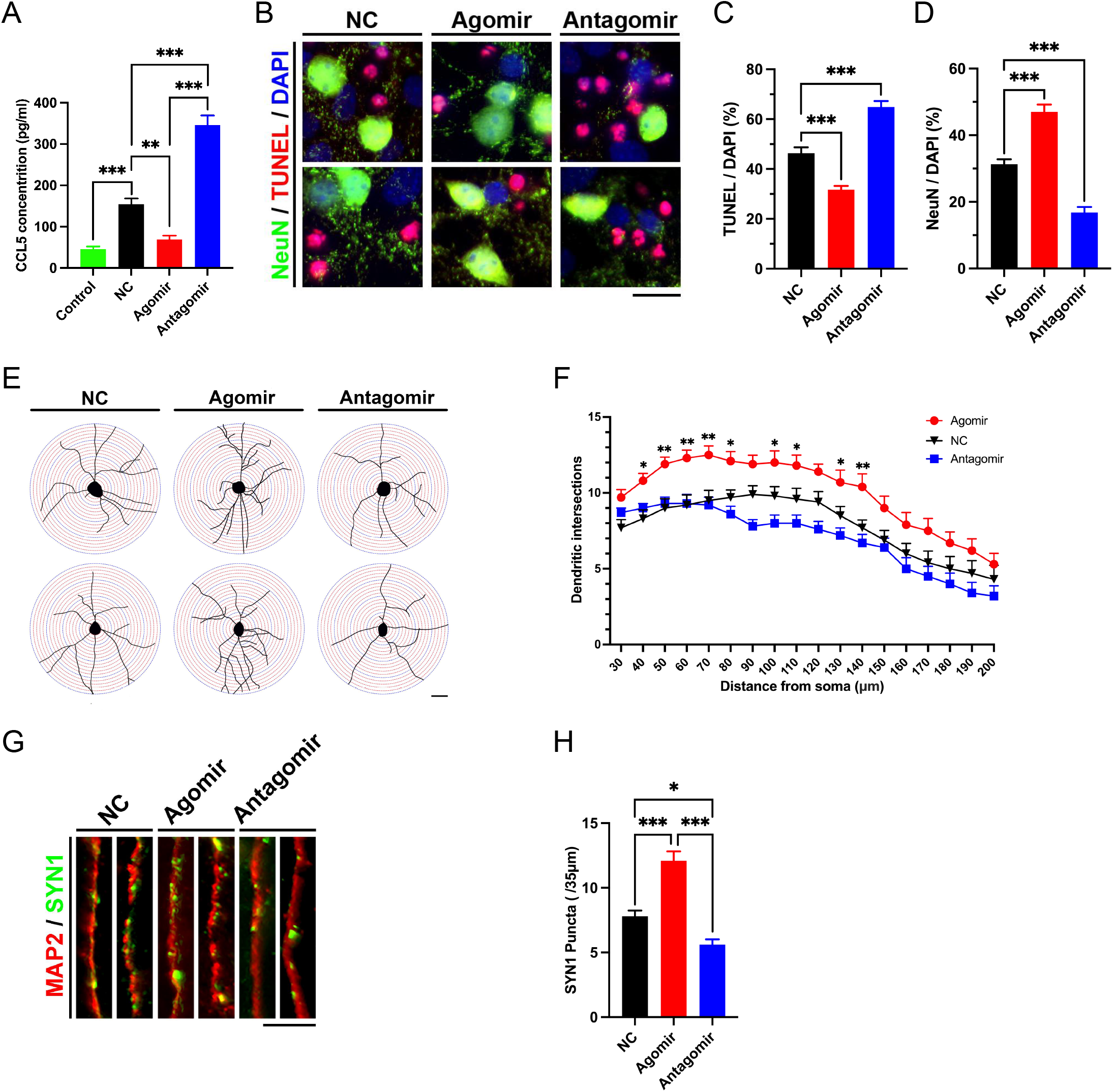
Overexpression of miR-324-5p protects cortical neurons from OGD injury in astrocyte-neuron co-cultures. (A) ELISA quantification of CCL5 concentrations in the medium of NC agomir-, miR-324-5p agomir-, and miR-324-5p antagomir-treated co-cultures, and in un-OGD-treated controls (*n*=5). (B) Representative immunofluorescence images of NeuN and TUNEL co-labeling. Scale bar: 20 μm. (C and D) Proportions of apoptotic cells (C) and NeuN-positive neurons (D) in NC agomir-, miR-324-5p agomir-, and miR-324-5p antagomir-treated co-cultures (*n*=10). (E and F) Representative tracings of MAP2-stained neurons (E) and Sholl analysis (F) of neuronal dendritic morphology in NC agomir-, miR-324-5p agomir-, and miR-324-5p antagomir-treated co-cultures (*n*=10). Scale bar: 50 μm. (G and H) Representative immunofluorescence images of MAP2 and SYN1 co-labeling (G) and synapse puncta density (H) in NC agomir-, miR-324-5p agomir-, and miR-324-5p antagomir-treated co-cultured neurons (*n*=10). Scale bar: 10 μm. Statistical comparisons in F were performed by two-way ANOVA with Tukey’s post-hoc test; all other panels by one-way ANOVA with Tukey’s post-hoc test.

TUNEL assay results indicated that at D3 post-OGD, apoptotic cells were significantly fewer in the agomir group compared with the NC groups (Figure 5B-C). Furthermore, the proportion of surviving NeuN-positive neurons was significantly higher in the agomir group than in the NC control (Figure 5D). In contrast, the antagomir group displayed more apoptotic cells than the NC group, and there were fewer NeuN-positive neurons observed in the antagomir group. These results demonstrate that upregulating miR-324-5p expression in the co-culture system reduces neuronal apoptosis after OGD, thereby exerting a protective effect on neuronal survival. Conversely, suppressing miR-324-5p expression exacerbates neuronal apoptosis following OGD injury.

Sholl analysis demonstrated that at D3 and D6 post-OGD, the dendrite intersections of neurons in the miR-324-5p agomir group were significantly more than those in the NC control group (Figure 5E-F, Supplementary Figure S3C). At D6 post-OGD, neurons in the agomir group exhibited significantly more dendritic branches within distances of 40-80 μm, 100-110 μm, and 130-140 μm from the soma compared with those in the NC group. No significant differences were detected between the antagomir and NC groups. These findings indicate that miR-324-5p upregulation significantly preserves neuronal dendritic structures after OGD injury, whereas its downregulation simplifies dendritic structure, potentially impeding functional recovery of neurons after OGD treatment.

MAP2/SYN1 staining results revealed that at D6 post-OGD, synaptic densities in the agomir group were significantly higher than in the NC group (Figure 5G-H). Conversely, neurons in the antagomir group exhibited significantly reduced synaptic densities compared with the NC group. These findings indicate that enhancing miR-324-5p expression in the co-culture system significantly preserves synaptic density post-OGD, while its inhibition leads to greater synaptic loss.

To assess the impact of regulating the miR-324-5p/CCL5 axis on intracellular signaling pathways in the cortex after ischemic injury, we collected total proteins from the IP and CP regions of MCAO mice injected with BSA, CCL5 antibody, rCCL5, or MVC. SDS-PAGE and Western blot analysis revealed that on D3 post-MCAO, the levels of activated ERK, as indicated by the phosphorylated ERK (p-ERK) to total ERK (t-ERK) ratio, were significantly lower in the IP regions compared with the CP regions in the BSA control group (Figure 6A-B). In the anti-CCL5 group, p-ERK ratios were significantly higher than in the BSA control group in the IP regions. Following MVC injection, a significant increase in the p-ERK ratio in the IP regions was observed. In the rCCL5 group, p-ERK ratio showed a significant decrease in the IP regions compared with the CP regions, with no statistical differences observed compared with the BSA control.

**Figure 6.**
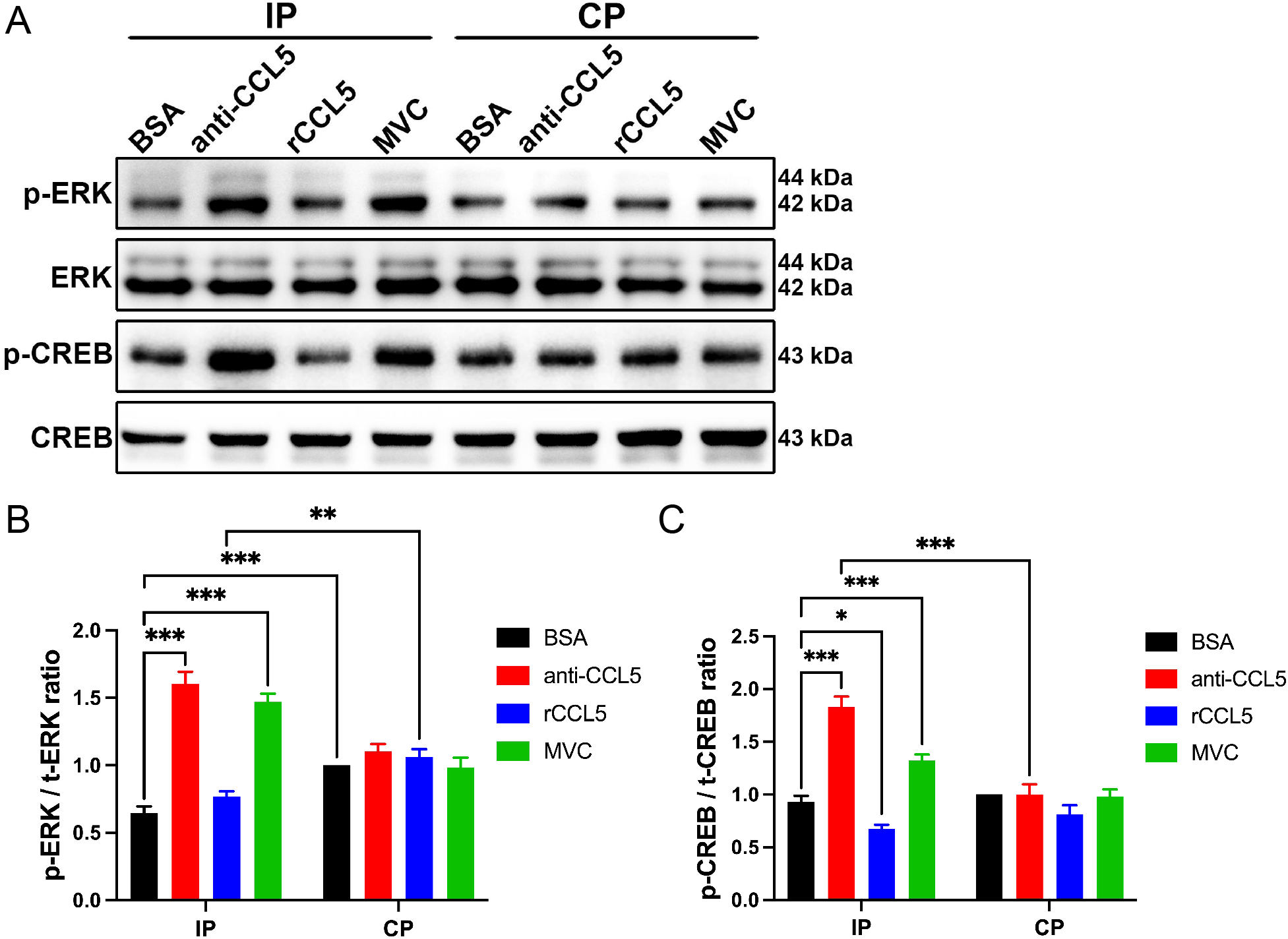
CCL5 antibody delivery and CCR5 blockade upregulate ERK/CREB signaling in the ipsilateral cortex of MCAO mice. (A) Representative Western blot images of p-ERK, t-ERK, p-CREB, and t-CREB in cortical lysates from the IP and CP regions of BSA-, CCL5 antibody-, rCCL5-, and MVC-treated mice on D3 after MCAO. (B and C) Ratios of p-ERK to t-ERK (B), and p-CREB to t-CREB (C) in the IP and CP regions of BSA-, CCL5 antibody-, rCCL5-, and MVC-treated mice on D3 after MCAO (*n*=6). Protein levels were normalized to the CP region of BSA-treated group. Statistical comparisons were performed by two-way ANOVA with Tukey’s post-hoc test.

In the BSA group, no significant differences in phosphorylated CREB (p-CREB) to total CREB (t-CREB) ratios were observed between IP regions and CP regions at D3 post-injury (Figure 6C). In the anti-CCL5 group, levels of activated CREB in the IP regions were significantly higher compared with both the BSA control group and the CP regions. Following MVC injection, significant upregulation of activated CREB levels was observed in the IP regions. In the rCCL5 group, p-CREB ratios were significantly lower in the IP regions compared with the BSA control group. These results indicate that the ERK pathway is downregulated in the ipsilateral cortex following ischemic stroke. Inhibiting CCL5 expression or blocking the CCR5 receptor could restore the ERK/CREB regulatory pathway in the ipsilateral cortex of MCAO mice, potentially exerting a protective effect on the structure and function of neurons following ischemic injury.

Given that brain tissue lysates contain proteins from multiple cell types, including astrocytes, neurons, microglia, and endothelial cells, we next isolated primary cortical neurons and co-cultured them with astrocyte-conditioned medium (ACM) to specifically examine how the astrocytic miR-324-5p/CCL5 axis regulates neuronal ERK/CREB signaling following ischemic injury. After OGD treatment, neurons were co-cultured with OGD-ACM for three days, followed by supplementation with BSA, rCCL5, CCL5 antibody, or a combination of MVC and rCCL5 in the neuronal culture media (Figure 7A). Neuronal proteins were collected, and changes in the expression of the ERK/CREB pathway were examined. As shown in Figure 7B-D, primary cortical neurons co-cultured with OGD-ACM exhibited significantly decreased levels of activated ERK and CREB after being treated with rCCL5 for 10-60 minutes at D3 post-OGD. Conversely, the addition of CCL5 antibody or the simultaneous addition of 5 nM MVC and rCCL5 led to increased levels of activated ERK and CREB in neurons. These results suggest that post-OGD, CCL5 inhibits the neuronal ERK/CREB pathway via the CCR5 receptor. Decreasing CCL5 expression or blocking the neuronal CCR5 receptor both restored the ERK/CREB pathway in neurons post-OGD injury.

**Figure 7.**
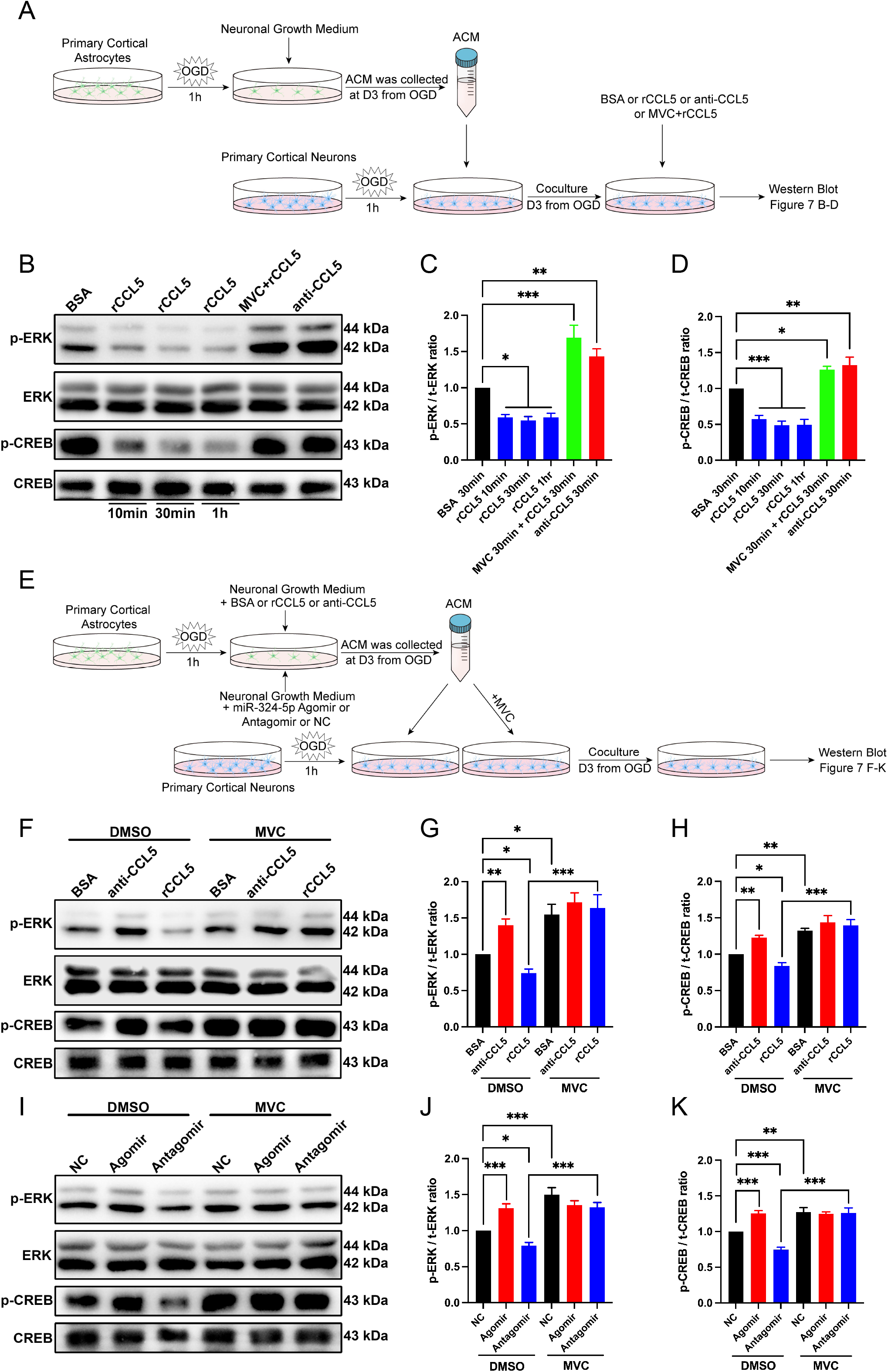
CCL5 inhibition and CCR5 blockade upregulate the neuronal ERK/CREB pathway after OGD injury. (A) Schematic of the ACM-neuron co-culture experimental design for panels B-D. (B) Representative Western blot images of p-ERK, t-ERK, p-CREB, and t-CREB in lysates from BSA-, CCL5 antibody-, rCCL5-, and MVC+rCCL5-treated neurons. (C and D) Ratios of p-ERK to t-ERK (C), and p-CREB to t-CREB (D) in BSA-, CCL5 antibody-, rCCL5-, and MVC+rCCL5-treated neuronal lysates (*n*=6). Protein levels were normalized to the BSA group. (E) Schematic of the ACM-neuron co-culture experimental design for panels F-K. (F) Representative Western blot images of p-ERK, t-ERK, p-CREB, and t-CREB in neurons co-cultured with BSA OGD-ACM, anti-CCL5 OGD-ACM, or rCCL5 OGD-ACM. (G and H) Ratios of p-ERK to t-ERK (G), and p-CREB to t-CREB (H) in neurons co-cultured with BSA OGD-ACM, anti-CCL5 OGD-ACM, or rCCL5 OGD-ACM (*n*=5). Protein levels were normalized to the BSA+DMSO group. (I) Representative Western blot images of p-ERK, t-ERK, p-CREB, and t-CREB in neurons co-cultured with NC agomir OGD-ACM, miR-324-5p agomir OGD-ACM, or miR-324-5p antagomir OGD-ACM. (J and K) Ratios of p-ERK to t-ERK (J), and p-CREB to t-CREB (K) in neurons co-cultured with NC agomir OGD-ACM, miR-324-5p agomir OGD-ACM, or miR-324-5p antagomir OGD-ACM (*n*=5). Protein levels were normalized to the NC agomir+DMSO group. Statistical comparisons in C and D were performed by one-way ANOVA with Dunnett’s post-hoc test; all other panels by one-way ANOVA with Tukey’s post-hoc test.

After OGD treatment of primary cortical astrocytes, the culture medium was supplemented with BSA, CCL5 antibody, or rCCL5. The respective OGD-ACMs (BSA OGD-ACM, anti-CCL5 OGD-ACM, and rCCL5 OGD-ACM) were collected, and OGD-treated neurons were co-cultured with these OGD-ACMs for three days (Figure 7E). Subsequent Western blot results showed significantly increased levels of activated ERK and CREB in neurons co-cultured with anti-CCL5 OGD-ACM compared with the BSA OGD-ACM control group (Figure 7F-H). Conversely, decreased levels of activated ERK and CREB were observed in neurons co-cultured with rCCL5 OGD-ACM.

Furthermore, when MVC was added to the ACM-neuron co-culture systems at a final concentration of 5 nM in each group immediately post-OGD, Western blot results revealed significantly upregulated ERK/CREB pathway activity in neurons co-cultured with both BSA OGD-ACM and rCCL5 OGD-ACM compared with the corresponding groups without MVC treatment. These results further indicate that both inhibiting CCL5 expression and blocking the CCR5 receptor could significantly enhance the ERK/CREB pathway in neurons post-OGD.

After OGD treatment, primary cortical astrocytes were transfected with NC agomir, miR-324-5p agomir, or miR-324-5p antagomir, respectively. The respective OGD-ACMs (NC OGD-ACM, agomir OGD-ACM, and antagomir OGD-ACM) were collected, and OGD-treated neurons were co-cultured with these OGD-ACMs for three days (Figure 7E). Western blot results showed significantly increased levels of activated ERK and CREB in neurons co-cultured with agomir OGD-ACM compared with the NC OGD-ACM group (Figure 7I-K). Additionally, decreased levels of activated ERK and CREB were detected in antagomir OGD-ACM co-cultured neurons.

Following MVC treatment, significant upregulation of the ERK/CREB pathway was observed in neurons co-cultured with NC OGD-ACM and antagomir OGD-ACM, compared with the corresponding groups without MVC treatment. Moreover, no statistical differences were detected in the activated ERK and CREB levels among neurons co-cultured with agomir OGD-ACM, either with or without MVC addition, and neurons co-cultured with NC OGD-ACM with MVC addition. This indicated that astrocytic miR-324-5p primarily affects the co-cultured neuronal ERK/CREB pathway through influencing the CCR5 receptor. Taken together, these results indicate that overexpressing miR-324-5p in astrocytes can significantly upregulate the neuronal ERK/CREB pathway via the CCR5 receptor post-OGD, whereas inhibiting astrocytic miR-324-5p decreased the neuronal ERK/CREB pathway (Figure 8).

**Figure 8.**
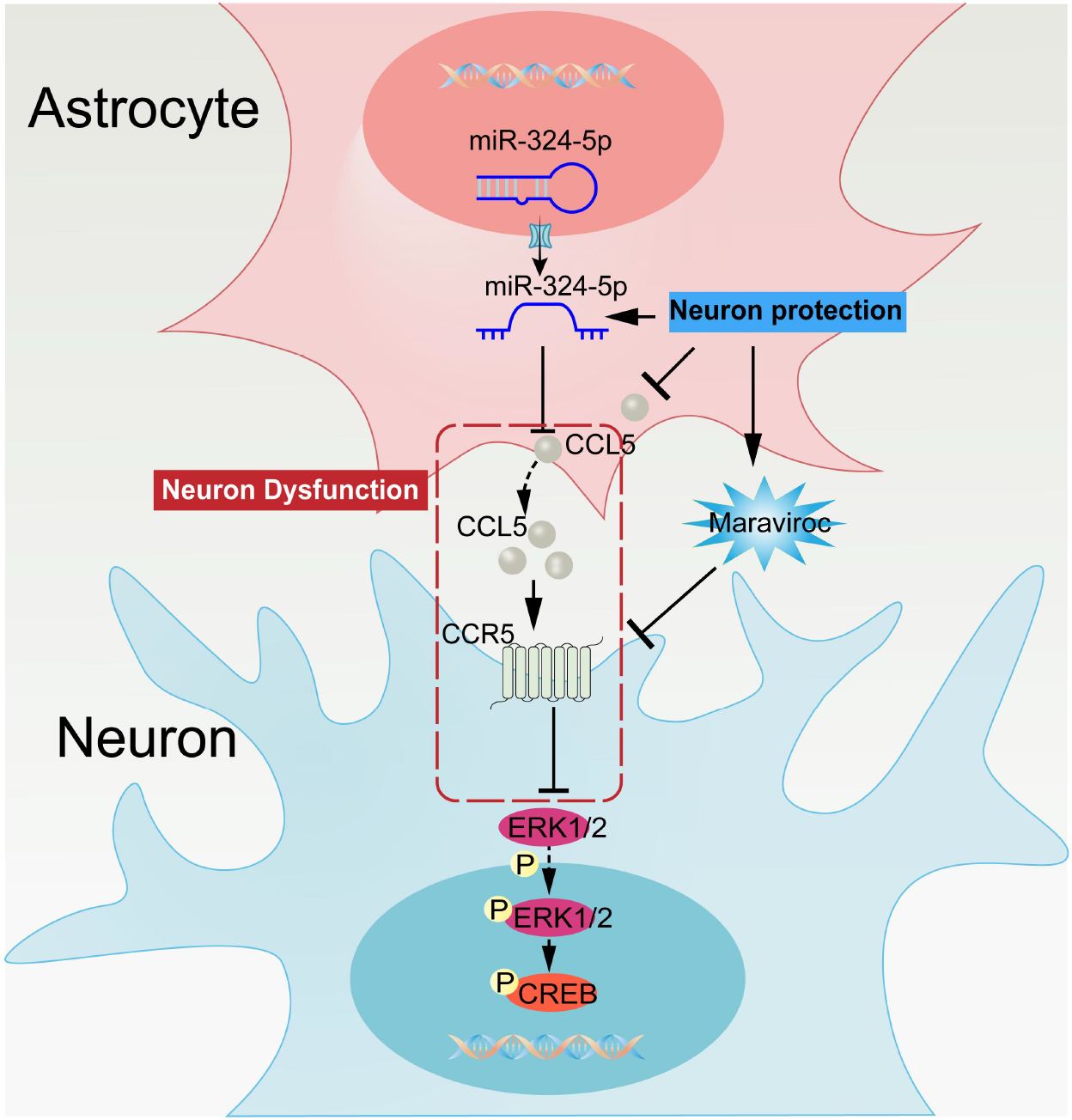
Proposed model of neuroprotection by astrocytic miR-324-5p through CCL5 inhibition and regulation of neuronal CCR5/ERK/CREB signaling after ischemic stroke.

## Discussion

The main findings of this study are as follows: (1) Following ischemic stroke, Ccl5 mRNA expression increased from D1 to D7, while miR-324-5p was significantly decreased after D1 in the ipsilateral cortex of MCAO mice; (2) Inhibiting CCL5 expression through intracerebral CCL5 antibody or miR-324-5p delivery attenuated astrocyte reactivity and microglial activation, reduced the infarct area, protected neurological function, and preserved both dendritic complexity and dendritic spine density in MCAO mice; (3) In the astrocyte-neuron co-culture system, inhibiting astrocytic CCL5 expression with CCL5 antibody or miR-324-5p following OGD decreased neuronal apoptosis and preserved dendritic complexity and synapse density; (4) Intracerebral CCL5 antibody delivery upregulated the ERK/CREB pathway in the ipsilateral cortex of MCAO mice; (5) Inhibiting astrocytic CCL5 expression by CCL5 antibody or miR-324-5p upregulated the ERK/CREB pathway in ACM-cocultured neurons after OGD treatment; (6) MVC application reduced infarct area in MCAO mice, decreased neuronal apoptosis, and preserved dendritic complexity and synapse density in OGD-treated neurons *in vitro*; (7) MVC application upregulated the ERK/CREB pathway in ACM-cocultured neurons after OGD treatment.

Our study reinforces the finding that CCL5 inhibition in experimental stroke models reduced infarct size, improved neurological function, suppressed glial reactivity, and preserved both dendritic field complexity and dendritic spine density after stroke. Ccl5 mRNA expression in the ipsilateral cortex was elevated from D1 to D7 after ischemic stroke (Figure 1C), and CCL5 protein concentrations in the IP region were correspondingly increased from D3 to D7 (Figure 2A). CCL5 inhibition resulted in reduced infarct size, improved neurological scores, and enhanced motor functions in MCAO mice (Figure 2B-E), while exogenous rCCL5 administration exacerbated infarct size (Figure 2B-C). Astrocyte surface area and microglial surface area were significantly reduced in CCL5-inhibited mice, whereas rCCL5 administration increased microglial surface area at D3 post-stroke (Figure 2G-J). Furthermore, CCL5 inhibition preserved dendritic field complexity and dendritic spine density, while rCCL5 administration reduced dendritic field complexity.

In our previous work, we demonstrated that astrocytic miR-324-5p suppresses Ccl5 mRNA expression (C. Sun et al. 2019). We therefore investigated whether overexpressing miR-324-5p in the ipsilateral cortex could recapitulate the effect of CCL5 inhibition in MCAO mice. Notably, FISH analysis revealed that miR-324-5p expression is significantly enriched in astrocytes relative to microglia in the peri-infarct region (Figure 1G), supporting the hypothesis that the miR-324-5p/CCL5 regulatory axis operates preferentially within astrocytes following ischemic stroke. Indeed, miR-324-5p agomir reduced CCL5 protein concentrations, while miR-324-5p antagomir elevated them in the ipsilateral cortex after ischemic stroke (Figure 3A). Administration of miR-324-5p agomir reduced infarct size, improved Longa scores, and enhanced motor functions, while miR-324-5p antagomir increased infarct size (Figure 3B-E). Both the CCL5 antibody and the miR-324-5p agomir reduced CCL5 expression in the ipsilateral cortex and improved motor functions in MCAO mice. The CCL5 antibody group exhibited notably better neurological scores through D3, although by D7, both groups showed comparable scores (Figures 2D, 3D). Moreover, the rotarod test revealed better motor performance in the CCL5 antibody group compared with the miR-324-5p agomir group at D3 and D7 post-MCAO (Figures 2E, 3E). These differences may be attributed to the immediate neutralization of extracellular CCL5 by the antibody, whereas the miRNA agomir requires time to suppress CCL5 expression post-transcriptionally following transfection. Therefore, reducing CCL5 levels early after stroke may confer greater benefits in enhancing motor recovery.

Decreased reactive astrocyte density and surface area were observed in miR-324-5p agomir-treated mice from D3 to D7 (Figure 3G-J). In contrast, miR-324-5p antagomir induced excessive astrocyte reactivity in the early stage post stroke (Figure 3G). Furthermore, miR-324-5p agomir maintained cortical microglia in a less-activated state during the early stage following stroke, as indicated by the reduced microglial surface area shown in Figure 3K.

The acute inflammatory response to stroke involves complex reactions to sterile tissue injury, which are essential for initiating processes that facilitate the clearance of damaged tissue and establish an optimal environment for subsequent tissue repair (Y. Mo et al. 2020). However, excessive inflammatory activation can lead to the overproduction of detrimental mediators, including cytokines, chemokines, and reactive oxygen species (ROS). This inflammatory cascade following stroke may compromise vascular integrity, promote cellular apoptosis, and contribute to secondary brain damage, thereby exacerbating cerebral lesions (Cheng et al. 2022).

Microglia, the resident immune cells of the brain, become rapidly activated following stroke, exhibiting both pro-inflammatory and anti-inflammatory roles at different stages of ischemic stroke (Xu et al. 2020). During the acute phase, activated microglia release a range of inflammatory mediators including TNF, IFN-β, and IL-6, contributing to a robust inflammatory response that impedes neural recovery (Lambertsen et al. 2012). In the subacute phase, microglia shift towards a protective role by secreting neurotrophic factors such as IGF1, which aids in neural repair and remodeling following ischemic injury (Lalancette-Hébert et al. 2007).

Our data revealed that both miR-324-5p overexpression and CCL5 inhibition reduced reactive astrocyte surface area and activated microglia surface area after stroke. The tight coupling of morphological features and functional states is well established in both astrocytes and microglia (Karperien et al. 2013; Gomez-Nicola and Perry 2015). Astrocytic reactivity is characterized by upregulation of GFAP and the extension of processes toward the infarct region, contributing to glial scar formation (Escartin et al. 2021). Microglial activation involves the transformation from a ramified to an amoeboid morphology (Davis et al. 2017). The reduction in glial surface area thus reflects a lower degree of glial reactivity in the peri-infarct cortex, suggesting an attenuation of the post-ischemic glial response. Compared with anti-CCL5 treatment, miR-324-5p agomir administration not only reduced reactive astrocyte surface area but also reduced reactive astrocyte density. This difference may reflect the ability of miR-324-5p agomir to continuously inhibit CCL5 expression throughout the experimental period, and/or its capacity to suppress other target genes that cooperate with CCL5 in driving astrocyte reactivity in the ischemic cortex.

It has been reported that the administration of exogenous RNA can induce activation of resident microglia in the cortex (Z. Chen et al. 2021). Microglia density did not change significantly in the anti-CCL5 group or in the agomir group compared with their respective controls (Supplementary Figures S1C-D, S2A-B). However, a direct comparison between these two groups showed that the CCL5 antibody was associated with greater reductions in microglial surface area at D7 than the miR-324-5p agomir group (Figures 2J, 3L). This difference was likely attributable to excessive microglial stimulation by the exogenous RNAs, and/or to off-target effects of miR-324-5p on other genes in the ischemic cortex. In summary, both the CCL5 antibody and miR-324-5p agomir attenuated glial reactivity following ischemic stroke. Of note, miR-324-5p agomir administration exhibited a more pronounced inhibitory effect on astrocyte reactivity, whereas CCL5 antibody administration more effectively suppressed microglia activation.

Numerous studies have highlighted the association between neural network reorganization in the peri-infarct cortex and functional improvement in both stroke patients and animal models of experimental stroke (Dunne et al. 2016; L. L. Kong et al. 2014; Rossi and Zlotnik 2000). On the contrary, progressive neurodegeneration combined with reductions in dendritic density and synaptic density may exacerbate brain damage and compromise neurological function in the subacute phase of brain injury (Wu et al. 2023). In the present study, injections of rCCL5 or miR-324-5p antagomir tended to simplify dendritic branch structure, although neither total spine density nor mushroom spine density reached statistical significance compared with the respective control groups at D7 post-stroke (Figures 2K-O, 3M-Q). Conversely, both miR-324-5p agomir and CCL5 antibody treatment significantly preserved dendritic field complexity and dendritic spine density after ischemic stroke. This preservation of neuronal structure likely contributed to the reduced neurological deficits and infarct size observed in these treatment groups.

Activation of CCR5 has been shown to inhibit hippocampal long-term potentiation and modulate MAPK signaling and CREB phosphorylation (Marciniak et al. 2015; M. Zhou et al. 2016). In an HIV-induced dementia mouse model, neuronal CCR5 downregulation enhanced cortical neuronal plasticity and facilitated recovery of learning and memory by increasing CREB phosphorylation (M. Zhou et al. 2016). Post-stroke knockdown of CCR5 in the premotor cortex facilitates motor control recovery, accompanied by dendritic spine preservation, new patterns of cortical projections to the contralateral premotor cortex, and enhanced CREB and DLK signaling (Joy et al. 2019). Moreover, a loss-of-function CCR5 mutation in stroke patients has been associated with improved recovery across distinct measures of cognitive, motor, and sensory function, including memory, verbal functioning, and attention (Joy et al. 2019). In addition, CCR5 blockade following stroke improved axon mapping and synaptic density, and attenuated reactive gliosis and peripheral immune cell infiltration (Wu et al. 2023). Furthermore, activated CCR5 has been reported to induce neutrophil migration through the Akt/GSK-3β pathway, aggravating neuroinflammation during the acute stroke phase (C. Chen et al. 2020). Accordingly, CCR5 blockade substantially reduced the infiltration of peripheral immune cells and suppressed the reactive proliferation of glial cells after stroke (Wu et al. 2023).

MVC, a CCR5 antagonist widely used in the clinic, has demonstrated a favorable safety profile and the capacity to penetrate the blood-brain barrier (Han et al. 2012). MVC inhibits CCR5 activation, thereby impeding inflammatory chemokine-mediated recruitment of immune cells and limiting local inflammation progression (Kraft-Terry et al. 2010; Escola et al. 2010). In ischemic stroke, MVC treatment has been shown to attenuate levels of pro-inflammatory mediators IL-1β, IL-6, and TNF-α, suggesting a role in limiting local neural injury post-ischemia (Wu et al. 2023). In addition, our findings demonstrate that MVC also reduces CCL5 expression in astrocytes following OGD (Figure 4B). As CCL5 is a pro-inflammatory mediator and its inhibition attenuated astrocyte reactivity and microglial activation in MCAO mice, MVC may exert part of its anti-inflammatory activity by reducing CCL5 secretion.

MVC has been reported to promote sustained motor recovery and reduce infarct volumes following MCAO (B. Chen et al. 2022; Villanueva 2019). These neuroprotective effects are partly mediated through inhibition of neuronal apoptosis, evidenced by increased Bcl-2 expression and decreased BAX expression after ischemic injury (B. Chen et al. 2022). Consistent with these previous studies, TUNEL staining confirmed that MVC treatment inhibited neuronal apoptosis after OGD (Figure 4C-E). Furthermore, MVC treatment in MCAO mice reduced infarct volume (Figures 2B-C), suggesting that CCR5 blockade confers a neuronal survival benefit during ischemic stroke.

Both neutralization of CCL5 by the CCL5 antibody and blockade of the CCR5 receptor by MVC preserved neuronal dendritic structure and synaptic density following OGD injury (Figure 4F-I). Although CCL5 can activate CCR1, CCR3, and CCR5, all of which are expressed on neurons, Sholl analysis and SYN1 quantification showed no statistical differences between these two groups. This suggests that astrocyte-secreted CCL5 predominantly signals through the CCR5 receptor to modulate dendrite morphology and synapse density in OGD-treated neurons. Notably, the addition of exogenous rCCL5 at 20 ng/mL did not further reduce dendritic complexity after OGD treatment, suggesting that endogenous astrocyte-secreted CCL5 after OGD may have already exerted near-maximal suppressive effects on neuronal dendritic structure.

Transfection with miR-324-5p agomir reduced CCL5 concentrations in the astrocyte-neuron co-culture system, whereas miR-324-5p antagomir increased them (Figure 5A). Administration of miR-324-5p agomir phenocopied the effects of the CCL5 antibody in inhibiting neuronal apoptosis and preserving dendritic arbor organization and synapse density (Figure 5B-H). In contrast, miR-324-5p antagomir exacerbated neuronal apoptosis and synapse loss after OGD injury. Taken together with the *in vivo* finding that miR-324-5p agomir administration reduced infarct volume and improved neurological function in MCAO mice, these results suggest that suppression of astrocytic CCL5 expression via miR-324-5p plays a neuroprotective role in the context of ischemic stroke injury.

In neurodevelopment, the ERK/MAPK pathway plays a critical role in various processes including neuronal proliferation, synaptic growth, gliogenesis, and cerebral cortex development (Iroegbu et al. 2021). Following ischemic stroke, activation of the ERK signaling pathway confers neuroprotection, enhances neuronal plasticity and migration, and thereby significantly contributes to neural remodeling and repair (Liu et al. 2018). In addition, the ERK/MAPK pathway plays a crucial role in modulating angiogenesis in brain microvascular endothelial cells, potentially facilitating neuroprotection and neurological repair post-stroke (Song et al. 2023).

Activation of the Gαi signaling axis downstream of CCR5 suppresses intracellular cAMP levels and inhibits CREB phosphorylation in neurons (M. Zhou et al. 2016; Clarkson et al. 2011; Frank et al. 2018). CREB plays a key role in enhancing neuronal excitability and long-term synaptic plasticity (Y. Zhou et al. 2009; Dong et al. 2006; Lopez de Armentia et al. 2007; Viosca et al. 2009). Peak CREB phosphorylation and CREB-induced gene expression occur within the first two days following stroke (Sugiura et al. 2004), although CREB activation persists in glial cells for weeks post-stroke, contributing to both neurogenesis and gliogenesis (D. Y. Zhu et al. 2004; Liang et al. 2016). Moreover, CREB overexpression enhances the remapping of injured somatosensory and motor circuits and facilitates the formation of new functional connections within these circuits post-stroke, consequently, increasing CREB activity in motor neurons enhances motor recovery, while blocking CREB signaling impairs stroke recovery (Caracciolo et al. 2018).

In the present study, the levels of p-ERK and p-CREB were increased in the ischemic cortex after CCL5 antibody or MVC treatment in MCAO mice (Figure 6A-C). To specifically examine the ERK/CREB pathway in neurons, primary cortical neurons were isolated and co-cultured with OGD-ACM following OGD treatment. Addition of rCCL5 downregulated p-ERK and p-CREB levels in post-OGD neurons, whereas pre-treatment with MVC prevented these inhibitory effects of rCCL5, resulting in elevated levels of neuronal p-ERK and p-CREB (Figure 7B-D, 7F-H). Addition of the CCL5 antibody also upregulated the ERK/CREB pathway in neurons co-cultured with OGD-ACM. Collectively, these findings suggest that astrocytes increase CCL5 release following OGD, and that the secreted CCL5 inhibits the neuronal ERK/CREB pathway through the CCR5 receptor, an effect that can be blocked by the CCR5 antagonist MVC. Consequently, reduction of astrocytic CCL5 upregulates the neuronal ERK/CREB pathway, potentially explaining the molecular mechanism underlying the neuroprotective effects of the CCL5 antibody observed in ischemic stroke models.

Furthermore, MVC treatment increased the ERK/CREB pathway activity in neurons co-cultured with miR-324-5p antagomir-treated OGD-ACM, but did not further elevate this pathway in neurons co-cultured with miR-324-5p agomir-treated OGD-ACM (Figure 7I-K). This indicates that astrocytic miR-324-5p modulates the neuronal ERK/CREB signaling pathway primarily by suppressing CCL5 expression and thereby reducing CCR5 engagement on neurons. In summary, the present findings demonstrate the neuroprotective capacity of astrocytic miR-324-5p in preserving neuronal structure and enhancing motor recovery post-stroke through inhibition of CCL5 expression and consequent upregulation of the neuronal ERK/CREB pathway via the CCR5 receptor (Figure 8).

Primary cortical astrocytes were isolated and purified using the shaking method, yielding approximately 88% GFAP-positive cells at passage one, as confirmed by immunofluorescence characterization of astrocyte, neuronal, oligodendrocyte, and microglial markers (Supplementary Figure S4A-B). While minor contamination by non-astrocytic cell types cannot be entirely excluded, the astrocyte-enriched co-culture system employed here was sufficient to demonstrate the role of astrocytic CCL5 in modulating neuronal survival and dendritic integrity. It is also worth noting that astrocytes cultured in serum-containing medium undergo well-documented reactive changes, acquiring a flat, polygonal morphology with sparse processes, which may not fully recapitulate their *in vivo* properties. More refined purification strategies, such as immunopanning, have been shown to yield astrocytes with greater purity and better-preserved *in vivo* characteristics (Foo et al. 2011). Future studies employing such improved culture systems, together with cell-type-specific *in vivo* manipulation of CCL5 and miR-324-5p through viral or transgenic approaches, would further consolidate the conclusions of the present study.

## Methods

### Ethics statement

All animal experiments were approved by the Ethical Committee of the Experimental Animal Center of Shandong Second Medical University (Approval No. 2020SDL077) and were conducted in accordance with institutional guidelines for the care and use of laboratory animals. All surgery was performed under isoflurane anesthesia, and every effort was made to minimize suffering.

### Animals

Male C57BL/6J mice (20–25 g, 8–10 weeks) were obtained from the Experimental Animal Center of Shandong Second Medical University and housed under specific pathogen-free conditions on a 12 h light/dark cycle, with *ad libitum* access to water and standard rodent chow.

### MCAO model

Transient focal ischemia was induced by intraluminal filament occlusion of the right middle cerebral artery. Briefly, mice were anesthetized with 1.5–2% isoflurane, and core body temperature was maintained at 37 ± 0.5 °C throughout surgery using a feedback-controlled heating pad. The right common carotid artery (CCA) and external carotid artery (ECA) were exposed through a midline cervical incision. Following ligation of the ECA, a silicon-coated monofilament (Jialing, 1800AAA) was inserted via the CCA into the internal carotid artery to a depth of approximately 18 mm beyond the carotid bifurcation, thereby occluding the origin of the MCA. After 2 h of occlusion, the filament was gently withdrawn to allow reperfusion. Sham-operated mice underwent identical surgical procedures without filament insertion. Neurological deficits were scored immediately after surgery using the five-point Longa scale (Longa et al. 1989); only mice with a score of 3 were included and randomly assigned to experimental groups.

### Drug administration

For stereotaxic intracerebral injections, mice were anesthetized with isoflurane and secured in a stereotaxic frame. A total volume of 2 µL of the indicated agent was delivered at 0.2 µL/min using a Hamilton microsyringe (Hamilton, 6546005) at coordinates 1.0 mm anterior to bregma, 1.0 mm lateral (right) to the sagittal suture, and 1.0–1.5 mm below the dural surface. The needle was held in place for 5 min after injection to prevent reflux. The following agents were administered: NC agomir (200 µM), miR-324-5p agomir (200 µM), or miR-324-5p antagomir (200 µM) (all from GenePharma); BSA (20 ng/µL); anti-CCL5 antibody (R&D, MAB478, 0.2 µg/µL); or recombinant mouse CCL5 (R&D, 478-MR, 20 ng/µL). For systemic treatment, MVC (MedChemExpress, HY-13004) was administered intraperitoneally at a concentration of 5 µM in a total volume of 1 mL sterile saline at 4, 24, and 48 h after MCAO.

### Tissue collection and processing

Mice were euthanized at 1, 3, or 7 days after MCAO. For biochemical assays (qRT-PCR, ELISA, and Western blotting), brains were rapidly removed and sectioned into 1 mm coronal slices using a brain matrix. The peri-infarct cortex—defined as the 1–2 mm rim of cortical tissue immediately surrounding the visibly pale and opaque infarct core—was dissected from the second and third coronal slices, beginning from the most rostral aspect of the cerebral cortex. For immunofluorescence, mice were transcardially perfused with phosphate-buffered saline (PBS) followed by 4% paraformaldehyde (PFA) in PBS. Brains were post-fixed in 4% PFA/PBS overnight at 4 °C, cryoprotected in 30% sucrose/PBS until sinking, and embedded in OCT compound (Sakura, 4583) before sectioning.

### QRT-PCR

Total RNA was isolated using RNAiso Plus (TaKaRa, 9109) according to the manufacturer’s instructions. For mRNA analysis, cDNA was synthesized with the PrimeScript RT Reagent Kit with gDNA Eraser (TaKaRa, RR047A), and quantitative PCR was performed with TB Green Premix Ex Taq II (TaKaRa, RR820A) on an Applied Biosystems 7500 Real-Time PCR System. For miRNA analysis, reverse transcription and quantitative PCR were performed using the Mir-X miRNA qRT-PCR TB Green Kit (TaKaRa, 638314) according to the manufacturer’s instructions. mRNA and miRNA levels were normalized to Gapdh and U6 snRNA, respectively.

Primer sequences were as follows: *Ccl5* forward, 5’-GGAGTATTTCTACACCAGCAGCAAG-3’, reverse, 5’-GGCTAGGACTAGAGCAAGCAATGAC-3’; *Gapdh*forward, 5’-GGTGAAGGTCGGTGTGAACG-3’, reverse, 5’-CTCGCTCCTGGAAGATGGTG-3’. The miR-324-5p-specific primer set was purchased from Genecopoeia (MmiRQP0412). All reactions were performed in triplicate, and relative expression was calculated using the 2^-ΔΔCT^ method (Livak and Schmittgen 2001).

### TTC staining and infarct volume quantification

Infarct volume was measured by TTC staining. Serial 1 mm coronal brain sections were incubated in 2% TTC solution (Solarbio, G3005) at 37 °C for 30 min and subsequently fixed in 4% PFA/PBS. Images were captured with a digital camera, and infarct volumes were quantified by an investigator blinded to group assignment using ImageJ. To correct for edema-induced hemispheric swelling, infarct volume was expressed as a percentage of the contralateral hemisphere volume: [(contralateral volume − ipsilateral non-infarcted volume) / contralateral volume] × 100%.

### Behavioral testing

The five-point Longa scale was used to score focal neurological deficits: 0, no deficit; 1, failure to fully extend the contralateral forepaw; 2, circling contralaterally; 3, falling contralaterally; 4, no spontaneous locomotion. For the accelerating rotarod test, mice were placed on a rotating cylinder that accelerated from 4 to 40 rpm over 5 min before being held at 40 rpm for a further 5 min. The latency to fall was recorded only for animals that actively walked on the rotating rod. Trials were terminated without recording the time if the animal either fell without attempting to walk or gripped the rod and rotated passively for two consecutive revolutions. Animals were trained for three consecutive days before MCAO surgery.

### Immunofluorescence staining

Cryosections (35 µm) or cultured cells grown on glass coverslips were fixed in 4% PFA/PBS for 10 min, then permeabilized and blocked for 1 h at room temperature in PBS containing 0.3% Triton X-100 and 3% normal donkey serum. The following primary antibodies were used: mouse anti-GFAP (Cell Signaling Technology, 3670); rabbit anti-Iba1 (Wako, 019-19741); rabbit anti-MAP2 (Synaptic Systems, 188002); mouse anti-SYN1 (Synaptic Systems, 106011BT); and mouse anti-NeuN (Merck Millipore, MAB377). Primary antibodies were applied overnight at 4 °C, followed by incubation with fluorophore-conjugated secondary antibodies (Jackson ImmunoResearch) for 1 h at room temperature. Nuclei were counterstained with DAPI (Roche, 10236276001). Images were acquired on a Leica SP5 confocal microscope or an Olympus BX53 microscope. For brain sections, imaging was performed in the viable peri-infarct cortex immediately adjacent to the infarct core, where cells retained intact morphology. The infarct core boundary was identified by the loss of cellular staining integrity.

For NeuN/TUNEL co-labeling, NeuN immunofluorescence was performed first, followed by TUNEL staining using the In Situ Cell Death Detection Kit (Roche, 12156792910) according to the manufacturer’s instructions. DAPI counterstaining was applied as the final step.

For combined FISH and immunofluorescence, GFAP or Iba1 immunolabeling was carried out first. *In situ* hybridization for Ccl5 mRNA or miR-324-5p was then performed using the RNASweAMI in situ hybridization fluorescence detection kit (Servicebio, GF002 and GF003) per the manufacturer’s protocol, with the proteinase K digestion step omitted to preserve the immunofluorescence signals from the preceding antibody labeling.

### Golgi–Cox staining and dendritic morphology analysis

Golgi–Cox impregnation was performed using the FD Rapid GolgiStain Kit (FD NeuroTechnologies, PK401) according to the manufacturer’s instructions. Brains were immersed in impregnation solution for 14 days in the dark and then transferred to Solution C for 72 h. Cryosections (160 µm) were cut on a cryostat and imaged under brightfield conditions using a Leica SP5 microscope. Imaging was restricted to the peri-infarct cortex immediately adjacent to the infarct core, where Golgi-impregnated neurons with intact dendritic arbors were selected for analysis. The corresponding contralateral cortex served as a control region.

For Sholl analysis, cortical pyramidal neurons were randomly selected from the peri-infarct and contralateral cortices. Neuronal arbors were traced in Adobe Photoshop, and dendritic intersections with concentric circles at 10 µm intervals from 30 to 150 µm from the soma were counted by investigators blinded to group assignment.

For spine density analysis, segments of basal dendrites (25-100 µm in length, ≥ 50 µm from the soma) were randomly selected from the same regions. Spines were classified as mushroom, stubby, or thin based on head diameter and neck length (Bourne and Harris 2008), and spine density was quantified by blinded investigators.

### Primary astrocyte–neuron co-culture and OGD

Primary cortical astrocytes and neurons were prepared from newborn (within 24 h) C57BL/6J mice as previously described (Beaudoin et al. 2012; C. Sun et al. 2019). P1 cortical astrocytes were plated onto poly-D-lysine-coated 12-well plates and cultured until they reached ∼90% confluence. Primary cortical neurons dissociated with papain (Worthington, LS003126) were seeded at 5 × 10^4^ cells/well onto the astrocyte beds. After 9 days of co-culture (DIV9), OGD was induced by transferring plates to an anaerobic chamber (1% O_2_, 5% CO_2_, 94% N_2_) in glucose-free, serum-free DMEM (Gibco, 11966025) for 1 h. Plates were then returned to normoxic conditions in neuronal maintenance medium supplemented with one of the following at the indicated final concentrations: BSA (20 ng/mL), anti-CCL5 antibody (200 ng/mL), recombinant mouse CCL5 (20 ng/mL), CCL5 (20 ng/mL) + MVC (5 nM), NC agomir (20 nM), miR-324-5p agomir (20 nM), or miR-324-5p antagomir (20 nM). Co-cultured coverslips were fixed in 4% PFA/PBS at 3 days or 6 days post-OGD for immunofluorescence analysis.

To prepare ACM, P1 primary astrocytes in T75 flasks were subjected to OGD for 1 h, after which the medium was replaced with neuronal maintenance medium (without B27 or glutamine) containing the indicated proteins or miRNAs as described above. ACM was collected at day 3 and day 6 post-OGD, with medium replaced after the day 3 collection. Both collections were pooled, centrifuged at 1,000 × g for 5 min to remove debris, and stored at −80 °C. Control ACM was collected from normoxic astrocytes without OGD.

For neuronal protein extraction, primary cortical neurons were seeded onto poly-D-lysine-coated 6-well plates at 3 × 10^5^ cells/well; cytosine arabinofuranoside (Ara-C, 4 µM; Sigma-Aldrich, C1768) was added from DIV4 to suppress glial proliferation. OGD was performed at DIV9 as described above, after which medium was replaced with a 1:1 mixture of fresh neuronal maintenance medium and ACM. Neuronal protein was harvested 3 days after OGD.

### SDS-PAGE and Western blotting

Brain tissue or cultured neurons were lysed in ice-cold RIPA buffer (Thermo Fisher Scientific, 89900) supplemented with Halt Protease and Phosphatase Inhibitor Cocktail (Thermo Fisher Scientific, 78441). Lysates were clarified by centrifugation at 12,000 × g for 15 min at 4 °C. Protein concentration was determined by the BCA protein assay (Pierce, 23225). Equal amounts of protein (25-35 µg) were resolved on 10% SDS-polyacrylamide gels and transferred to PVDF membranes (Merck Millipore, ISEQ00010) using a wet-transfer system. Membranes were blocked with 5% BSA in TBST for 1 h at room temperature and then incubated overnight at 4 °C with the following primary antibodies: anti-phospho-ERK1/2 (Thr202/Tyr204) (Abcam, ab76299); anti-total ERK1/2 (Abcam, ab184699); anti-phospho-CREB (Ser133) (Abcam, ab32096); and anti-total CREB (Cell Signaling Technology, 9197). After incubation with HRP-conjugated secondary antibodies (Cell Signaling Technology) for 1 h at room temperature, signals were detected with SuperSignal West Pico PLUS chemiluminescent substrate (Thermo Fisher Scientific, 34580) and imaged on a Clinx ChemiScope 6000 imaging system. Band intensities were quantified by densitometry using ImageJ.

## Statistical analysis

All data are presented as mean ± SEM. Statistical analyses were performed using GraphPad Prism. For datasets with a single grouping factor, differences were assessed by one-way ANOVA followed by Tukey’s post hoc test for multiple comparisons among all groups, or Dunnett’s post hoc test when multiple experimental groups were compared against a single control group. For datasets with two grouping factors or repeated measures, two-way ANOVA followed by Tukey’s post hoc test was used. Comparisons between two groups were made using unpaired Student’s *t*-test where indicated. A *P* value < 0.05 was considered statistically significant. In all figures, statistical significance is denoted as \**p* < 0.05, \*\**p* < 0.01, and \*\*\**p* < 0.001.

## Acknowledgments

This work was supported by the National Natural Science Foundation of China (82001325 and 82070856), the Natural Science Foundation of Shandong Province (ZR2025MS1418 and ZR2020MH146), the Visiting Scholar Foundation of Shandong Province. We thank the Experimental Animal Center of Shandong Second Medical University for their kind assistance with the animal experiments.

---

**Figure S1.**
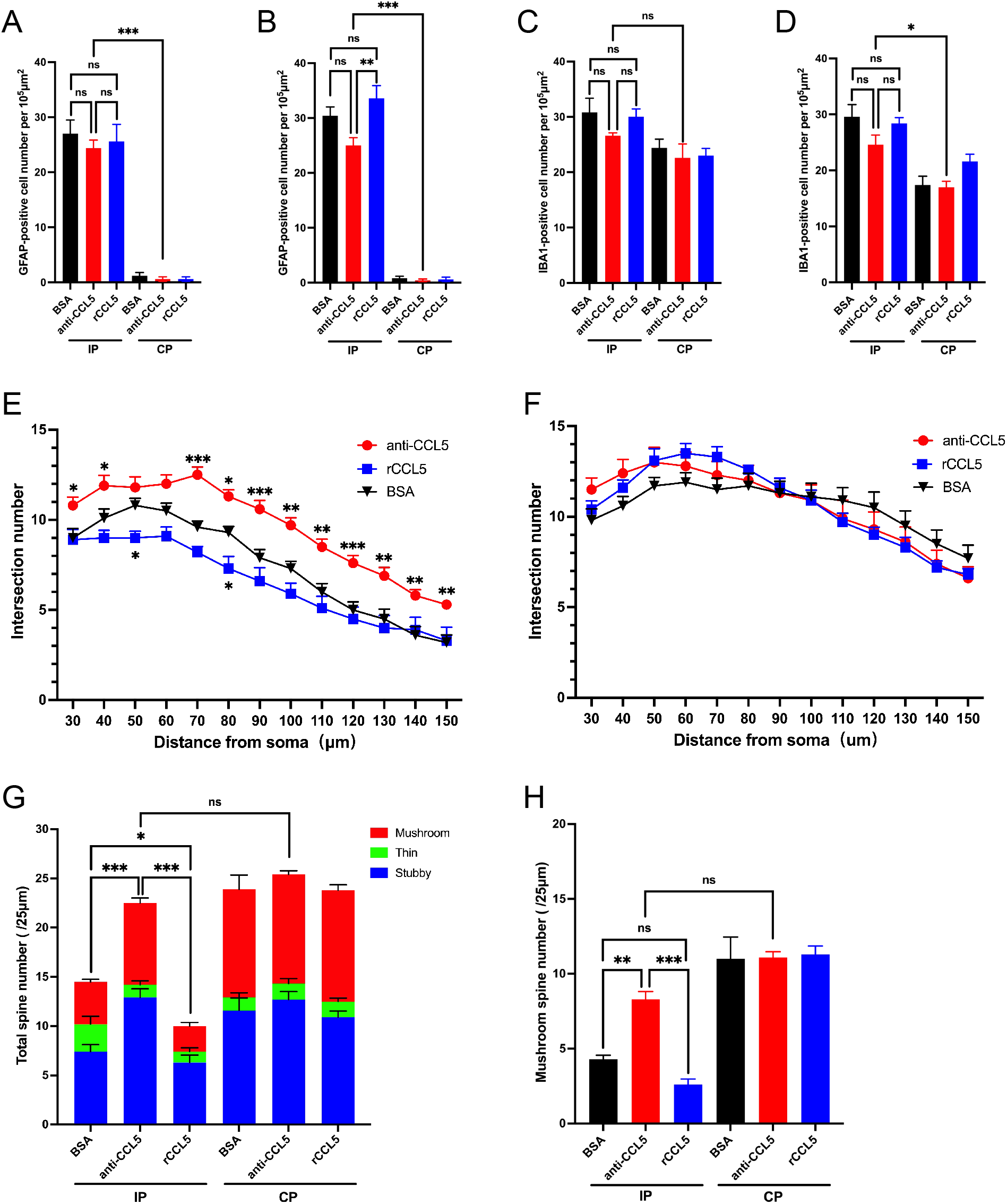
(A and B) Density of GFAP-positive astrocytes (cells per 10 µm²) in the IP and CP regions of BSA-, CCL5 antibody-, and rCCL5-treated mice on D3 (A) and D7 (B) after MCAO (*n*=5). (C and D) Density of IBA1-positive microglia (cells per 10 µm²) in the IP and CP regions of BSA-, CCL5 antibody-, and rCCL5-treated mice on D3 (C) and D7 (D) after MCAO (*n*=6). (E) Sholl analysis of dendritic branch complexity in the IP region of BSA-, CCL5 antibody-, and rCCL5-treated mice on D7 after MCAO (*n*=10). (F) Sholl analysis of dendritic branch complexity in the CP region of BSA-, CCL5 antibody-, and rCCL5-treated mice on D7 after MCAO (*n*=10). (G and H) Total spine density (G) and mushroom spine density (H) in the IP and CP regions of BSA-, CCL5 antibody-, and rCCL5-treated mice on D3 after MCAO (*n*=10). Stacked columns represent total spine density, with colored segments indicating the proportions of mushroom, stubby, and thin spines. Statistical comparisons in A-D, G, and H were performed by one-way ANOVA with Tukey’s post-hoc test. In E, statistical comparisons were performed by two-way ANOVA with Tukey’s post-hoc test.

**Figure S2.**
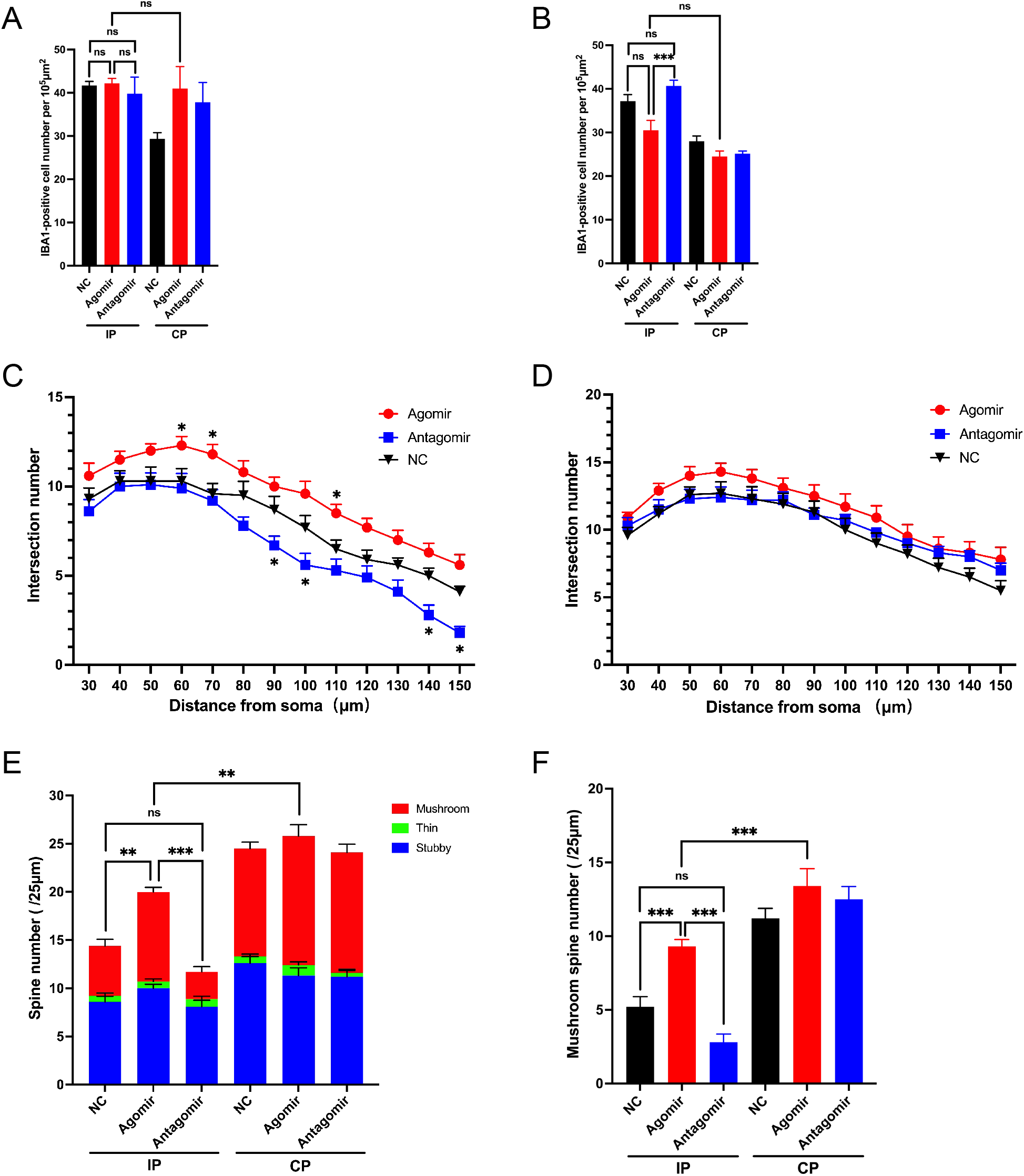
(A and B) Density of IBA1-positive microglia (cells per 10 µm²) in the IP and CP regions of NC agomir-, miR-324-5p agomir-, and miR-324-5p antagomir-treated mice on D3 (A) and D7 (B) after MCAO (*n*=5 in A, *n*=6 in B). (C) Sholl analysis of dendritic branch complexity in the IP region of NC agomir-, miR-324-5p agomir-, and miR-324-5p antagomir-treated mice on D7 after MCAO (*n*=10). (D) Sholl analysis of dendritic branch complexity in the CP region of NC agomir-, miR-324-5p agomir-, and miR-324-5p antagomir-treated mice on D7 after MCAO (*n*=10). (E and F) Total spine density (E) and mushroom spine density (F) in the IP and CP regions of NC agomir-, miR-324-5p agomir-, and miR-324-5p antagomir-treated mice on D3 after MCAO (*n*=10). Stacked columns represent total spine density, with colored segments indicating the proportions of mushroom, stubby, and thin spines. Statistical comparisons in A, B, E, and F were performed by one-way ANOVA with Tukey’s post-hoc test. In C, statistical comparisons were performed by two-way ANOVA with Tukey’s post-hoc test.

**Figure S3.**
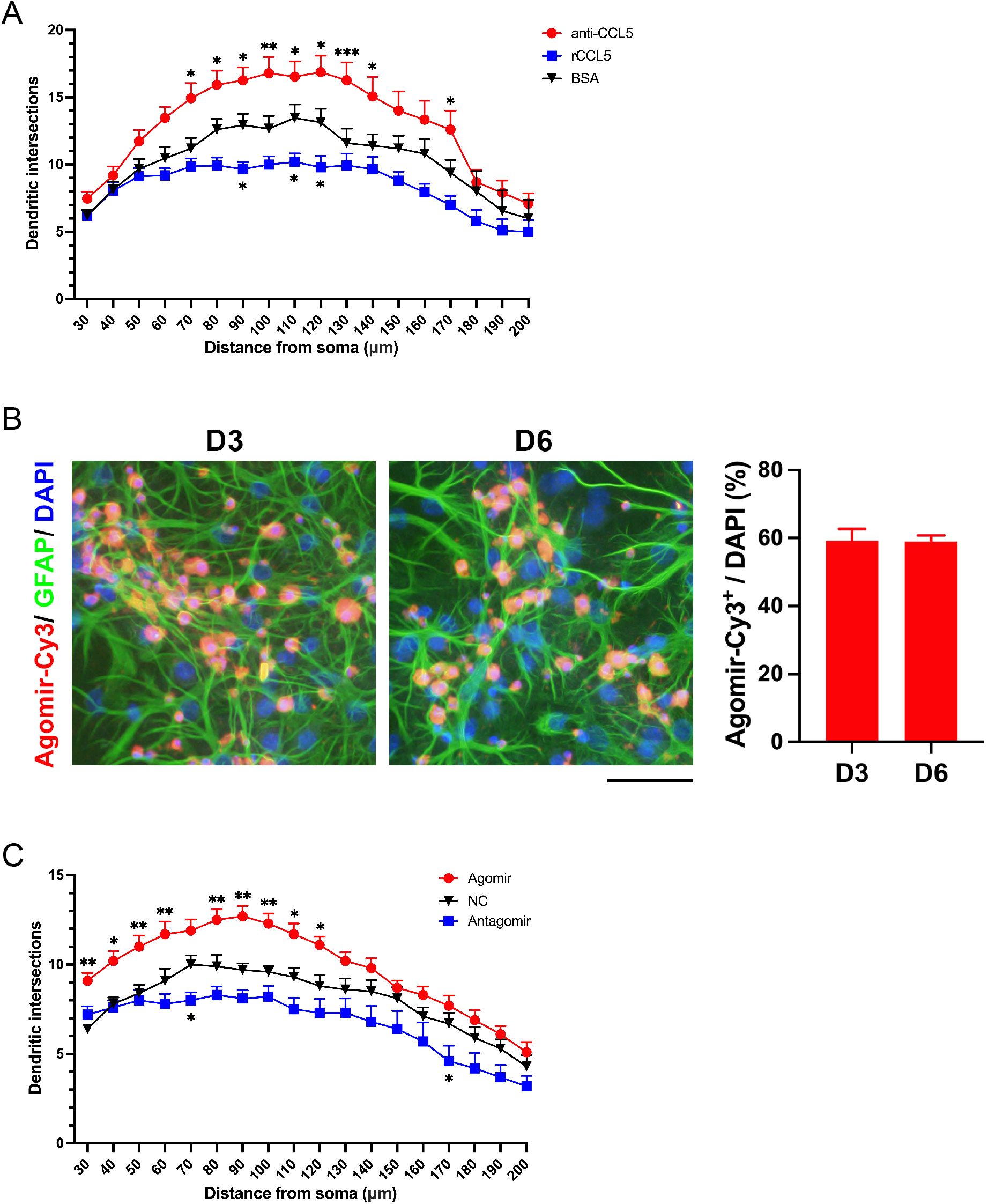
(A) Sholl analysis of dendritic branch complexity in BSA-, CCL5 antibody-, and rCCL5-treated co-cultured neurons on D3 after OGD (*n*=15). (B) Representative immunofluorescence images of GFAP labeling in astrocyte-neuron co-cultures following agomir-Cy3 transfection (left panel) and agomir transfection rate at D3 and D6 in co-cultured cells (right panel). (C) Sholl analysis of dendritic branch complexity in NC agomir-, miR-324-5p agomir-, and miR-324-5p antagomir-treated co-cultured neurons on D3 after OGD (*n*=10). Statistical comparisons in A and C were performed by two-way ANOVA with Tukey’s post-hoc test.

**Figure S4.**
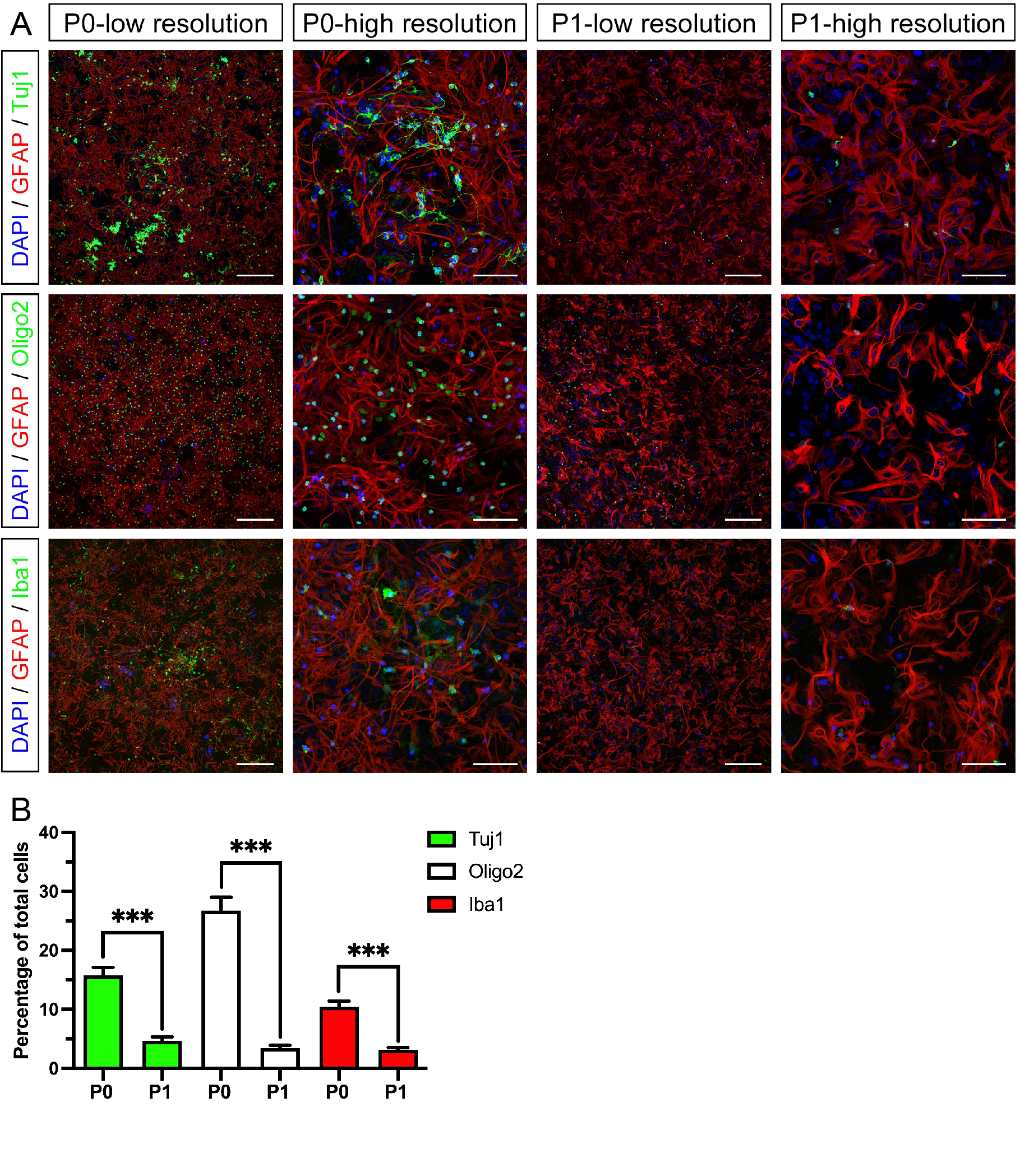
Characterization of primary cortical astrocyte purity. (A) Representative immunofluorescence images of GFAP co-labeled with Tuj1, Olig2, or Iba1 in P0 and P1 primary cortical astrocytes in vitro. Scale bar: 250 µm (low magnification) and 75 µm (high magnification). (B) Proportions of Tuj1-positive, Olig2-positive, and Iba1-positive cells among total DAPI-positive cells in P0 and P1 astrocyte cultures (*n*=6). Statistical comparisons between P0 and P1 were performed by unpaired t-test.

